# *Giardia intestinalis* deoxyadenosine kinase has a unique tetrameric structure that enables high substrate affinity and makes the parasite sensitive to deoxyadenosine analogues

**DOI:** 10.1101/2023.12.18.572228

**Authors:** Farahnaz Ranjbarian, Karim Rafie, Kasturika Shankar, Sascha Krakovka, Staffan G. Svärd, Lars-Anders Carlson, Anders Hofer

## Abstract

*Giardia intestinalis* is a protozoan parasite causing giardiasis, a severe, sometimes even life-threatening, diarrheal disease. *Giardia* is one of only a few known organisms that lack *de novo* synthesis of DNA building blocks, and the parasite is therefore completely dependent on salvaging deoxyribonucleosides from the host. The deoxyribonucleoside kinases (dNKs) needed for this salvage are generally divided into two structurally distinct families, thymidine kinase 1 (TK1)-like dNKs and non-TK1-like dNKs. We have characterized the *G. intestinalis* deoxyadenosine kinase and found that it, in contrast to previously studied non-TK1-like dNKs, has a tetrameric structure. Deoxyadenosine was the best natural substrate of the enzyme (K_M_=1.12 μM; V_max_=10.3 μmol·min^-1^·mg^-1^), whereas the affinities for deoxyguanosine, deoxyinosine and deoxycytidine were 400-2000 times lower. Deoxyadenosine analogues halogenated at the 2- and/or 2’
s-positions were also potent substrates, with comparable EC_50_ values as the main drug used today, metronidazole, but with the advantage of being usable on metronidazole-resistant parasites. Cryo-EM and 2.1 Å X-ray structures of the enzyme in complex with the product dAMP (and dADP) showed that the tetramer is kept together by extended N- and C-termini that reach across from one canonical dimer to the next in a novel dimer-dimer interaction. Removal of the two termini resulted in lost ability to form tetramers and a 100-fold decreased deoxyribonucleoside substrate affinity. This is the first example of a non-TK1-like dNK that has a higher substrate affinity as the result of a higher oligomeric state. The development of high substrate affinity could be an evolutionary key factor behind the ability of the parasite to survive solely on deoxyribonucleoside salvage.

**Authors summary:** The human pathogen *Giardia intestinalis* is one of only a few organisms that lack ribonucleotide reductase and is therefore completely dependent on salvaging deoxyribonucleosides from the host for the supply of DNA building blocks. We have characterized one of the *G. intestinalis* salvage enzymes, which was named deoxyadenosine kinase based on its substrate specificity. The enzyme also phosphorylated many deoxyadenosine analogues that were equally efficient in preventing parasite growth as the most used drug today, metronidazole, and also usable against metronidazole-resistant parasites. Structural analysis of the enzyme with cryo-EM and X-ray crystallography showed that the enzyme was unique in its family of deoxyribonucleoside kinases by forming a tetramer and mutational analysis showed that tetramerization is a prerequisite for the high substrate affinity of the enzyme. The ability to gain substrate affinity by increasing the number of enzyme subunits could potentially represent an evolutionary pathway that has assisted the parasite to become able to survive entirely on salvage synthesis of DNA building blocks.

---

The protozoan parasite *G. intestinalis* (synonymous to *G. lamblia* and *G. duodenalis*) causes giardiasis, which is an acute or chronic diarrheal disease spread by contaminated water or food and leads to 190 million symptomatic cases per year [1, 2]. However, the disease is also associated with a wide range of other symptoms including stomach cramps, vomiting, nausea, flatulence, dehydration and malabsorption, arthritis, and chronic fatigue syndrome. It can also start out as an asymptomatic infection but still cause malabsorption in children, followed by growth retardation, suppressed cognitive development and sometimes death. Treatment regimes are mainly based on metronidazole and to a lesser extent other 5-nitroimidazoles such as tinidazole. Metronidazole is associated with side effects such as nausea, abdominal pain, diarrhea, and in some cases neurotoxicity reactions [3]. Drug resistance to metronidazole and other 5-nitroimidazoles is emerging [4], highlighting the need to develop new drugs.

*G. intestinalis* is a diplomonad, which is a group of flagellated organisms characterized by having a duplicated cell content with two diploid nuclei in each cell [1]. The parasite has two life cycle stages, trophozoites and cysts. The trophozoite is the replicating form and attaches to the surface of epithelial cells in the duodenum where it proliferates and lives on the ingested food of the host. The trophozoites can also secrete a cyst-wall and go through two rounds of DNA replication to form cysts, which contain four nuclei and a 16N genome per cell (4N in each nucleus). The cysts, which leave the body in the feces, have only limited metabolic activity and are very resistant to diverse environmental factors. This makes them well adapted for survival outside the body for extended periods. When ingested, a combination of the acid pH in the stomach and the subsequent exposure to trypsin and bile in the duodenum induce parasite excystation into the replicating trophozoite form [5].

It is common that protozoan parasites lack *de novo* purine biosynthesis, which makes them sensitive to drugs targeting their compensatory salvage pathways needed to take up and use purines from the surroundings [6-9]. The drugs can either be inhibitors of the salvage process or more commonly act as substrate analogues. In the latter case, the salvage enzymes convert them to their nucleotide form and inhibit downstream processes such as nucleic acid synthesis, energy metabolism or other nucleotide-dependent reactions. In the case of *G. intestinalis*, it is not only *de novo* purine biosynthesis that is lacking. In fact, the parasite lacks *de novo* synthesis of all nucleotides including purines, pyrimidines, deoxyribonucleotides and thymidylate [10-12]. It is particularly striking that it lacks ribonucleotide reductase, which is a key enzyme for *de novo* synthesis of deoxyribonucleoside triphosphates (dNTPs) needed as building blocks for DNA [13]. Only very few organisms lack this enzyme, including *Giardia, Ureaplasma* as well as some species of *Entamoeba* and *Borrelia* [12, 14-16] The lack of ribonucleotide reductase makes *G. intestinalis* and other *Giardia* species completely dependent on salvaging deoxyribonucleosides from the surroundings to make the corresponding dNTPs. The first step in the salvage process is to phosphorylate the deoxyribonucleosides to the monophosphate form. The reaction is catalyzed by deoxyribonucleoside kinases (dNKs), which are considered rate-limiting in the salvage process [17]. The dNKs are subdivided into two structurally unrelated families, the thymidine kinase 1 (TK1) family and a family containing the remaining enzymes referred to as non-TK1-like dNKs [18]. The TK1 family has a strict substrate specificity adapted to phosphorylate thymidine-like substrates (thymidine and deoxyuridine) and serve the purpose to supply the cell with dTTP [18]. The thymidine kinase from *G. intestinalis* was recently characterized and this TK1-like enzyme was found to have a ∼10-fold higher affinity to thymidine than the human TK1 [19]. This could be an adaptation to be able to efficiently compete with the host cells and compensate for the lack of *de novo* dTTP synthesis in *G. intestinalis*. The high affinity makes the parasite sensitive to the substrate analogue azidothymidine, which is an efficient antigiardial agent in *in vitro* cell proliferation assays as well as in *G. intestinalis*-infected gerbils [19]. The incentive behind the current study was to investigate if also other dNKs in *G. intestinalis* have developed higher substrate affinities than the host enzymes as an adaptation to survive in the mammalian body. Studies of *G. intestinalis, T. brucei* and mammalian TK1 members have shown that substrate affinity can be gained by increasing the number of units the enzyme is composed [7, 19-21]. However, it has yet not been demonstrated for the non-TK1-like dNKs, which are in most cases dimers [17].

The non-TK1-like family of dNKs generally have much broader substrate specificities than the TK1 family. Mammalian cells have a full capacity to phosphorylate deoxyribonucleosides in the cytosol as well as the mitochondria and the single cytosolic member of this family, deoxycytidine kinase (dCK), is able to phosphorylate deoxycytidine as well as both purines [17]. In contrast, *G. intestinalis* lacks mitochondria and only needs cytosolic dNKs. Besides the thymidine kinase mentioned above, the parasite also has two non-TK1-like dNKs. Very little is known about them. A previous study showed that the dNK activities from a *G. intestinalis* extract separated as two peaks during purification, one for purine substrates and one for pyrimidines [22]. This does not match the current knowledge that there are three dNKs in total whereof one completely specific for thymidine. The lack of knowledge about these essential enzymes in the parasite has hampered the understanding of *Giardia* deoxyribonucleoside metabolism and hence its exploitation as a target for antiparasitic drugs.

In the current work, we have characterized one of the two non-TK1-like dNKs from *G. intestinalis*. The enzyme was found to use deoxyadenosine as its main substrate and was consequently given the name deoxyadenosine kinase (dAK). Several analogues of deoxyadenosine were also recognized as substrates, and cell studies showed that these analogues seem to be promising as drug candidates against the parasite. Structural analysis showed that *G. intestinalis* dAK is a homotetramer, which dramatically loses substrate affinity when tetramer formation is prevented by removal of the extended N- and C-termini at the dimer-dimer interface. For an organism that solely survives on deoxyribonucleosides it is of utmost importance to efficiently salvage them from the surroundings in the host. Increasing the number of units that the dNKs consist of may have been a fast-track evolutionary pathway of the *Giardia* genus to dramatically increase the substrate affinity and make itself independent of *de novo* dNTP synthesis.

## Results

### Sequence analysis of *G. intestinalis* dAK reveals N-terminal and C-terminal extensions

An alignment of the *G. intestinalis* dAK (ORF GL50803-17451 in assemblage A isolate WB) with several bacterial and eukaryotic non-TK1-like dNKs showed that it has an N-terminal extension that the other members lack (S1 Fig). Also, the C-terminus seems longer although the degree of homology between the species was not high enough to completely rule out that it was an alignment artefact at this stage of the analysis. The amino acid sequence has 40% identity to the second *G. intestinalis* non-TK1-like dNK and 36% identity to the *Mycoplasma mycoides* dAK, indicating a similarity to bacterial non-TK1-like dNKs [23]. For comparison, the corresponding identity to the metazoan dNKs, included in the sequence alignment, was only 24-29%. Conserved active site residues in the other species were present in the *G. intestinalis* dAK as well (S1 Fig).

### *G. intestinalis* dAK shows a preference for deoxyadenosine as a substrate and is inhibited by dATP

The *G. intestinalis* dAK gene was cloned into a pET-Z vector for expression in *Escherichia coli* as a fusion construct with a removable N-terminally His_6_-tagged Z protein followed by a tobacco etch virus (TEV) protease cleavage site. The expressed protein was purified by Nickel-NTA agarose chromatography, cleaved with TEV protease, and repurified with Nickel-NTA agarose to collect pure untagged dAK in the flow-through (Fig 1A). Based on initial enzyme activity studies, it was confirmed that the assay did not have any specific requirements regarding salts or reducing agents, and that it was linear with respect to time (S2 Fig). ATP was the best phosphate donor and deoxyadenosine the best substrate (Fig 1B-C). A kinetic analysis with deoxyadenosine as substrate showed that there was no obvious sign of cooperativity and that the enzyme had K_M_ and V_max_ values of 1.12 ± 0.37 μM and 10.3 ± 1.0 μmol·min^-1^·mg^-1^, respectively (Fig. 1D). It has thus a nearly 40-fold higher catalytic efficiency (k_cat_/K_M_) for deoxyadenosine than *M. mycoides* dAK [24].

**Fig 1.**
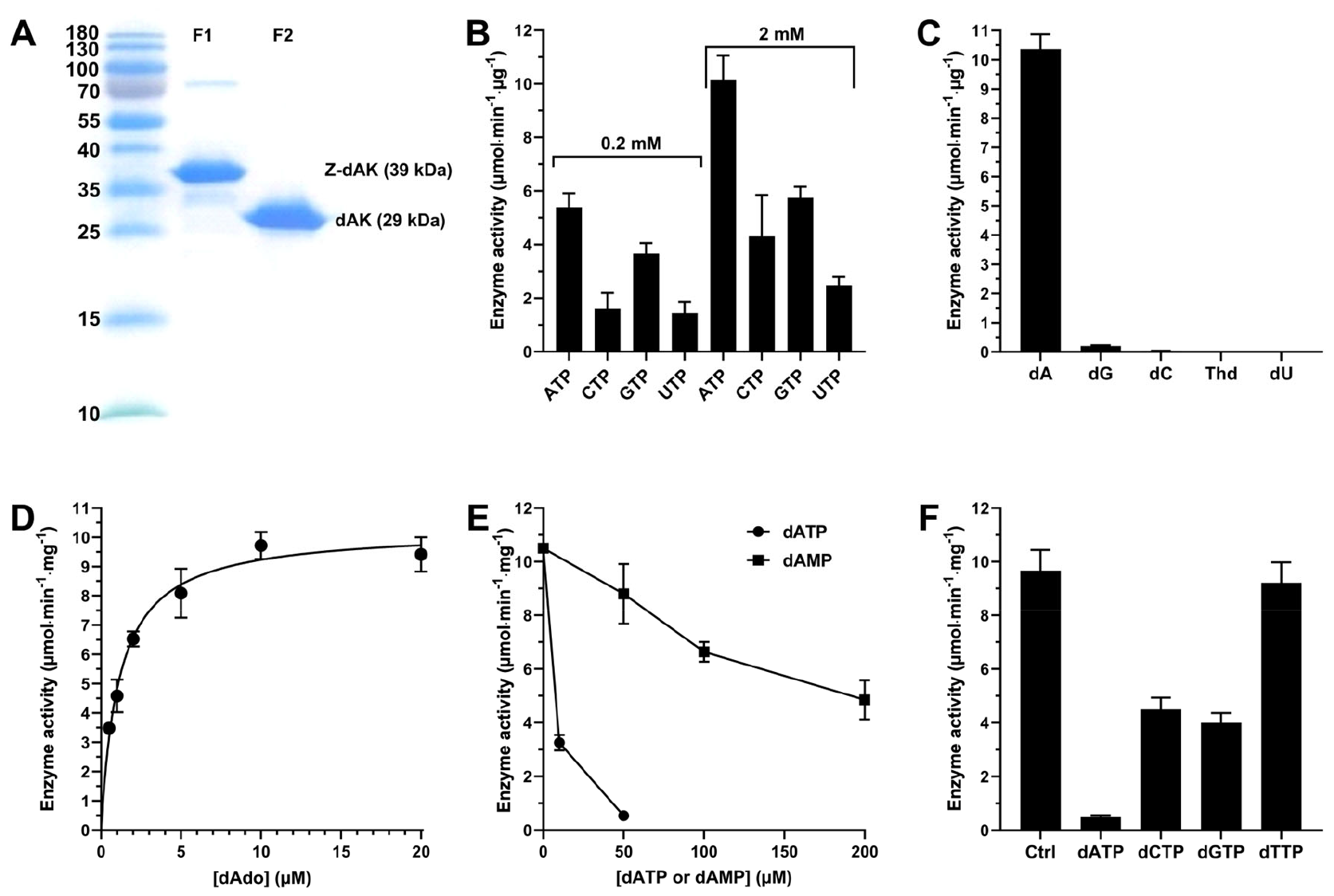
*G. intestinalis* dAK purification and characterization. (A) SDS-PAGE showing the purity of the protein after nickel agarose (F1), and after fusion partner removal and re-purification (F2). (B) Enzyme activity with different phosphate donors present at a concentration of 0.2 mM (left) or 2 mM (right). (C) Enzyme activity with all indicated substrates present simultaneously at a concentration of 200 μM in the assay. (D) Enzyme kinetic analysis with deoxyadenosine as substrate. (E) Enzyme inhibition with dATP and dAMP. (F) Effect of different dNTPs on enzyme activity. Standard enzyme assay conditions were used in the experiments and unless otherwise stated, the substrate was 200 μM deoxyadenosine and the phosphate donor 2 mM ATP. Graphs B-F represent the average of three independent experiments with standard errors indicated.

*G. intestinalis* dAK could be inhibited by its immediate product, dAMP, and with higher affinity by its far-end product, dATP (Fig. 1E). Further analysis showed that dATP was a mixed type of inhibitor with a K_i_ of 0.34 ± 0.05 μM (S3 Fig). Structural studies of other dNKs have demonstrated that the far-end dNTP product gives a dual inhibition where the deoxyribonucleoside moiety competes with the substrate and the phosphate groups mimic those of ATP but coming from the opposite direction [18]. The competition with ATP gives an effect of the V_max_ value and explains why the inhibition is mixed rather than competitive in assays where the deoxyribonucleoside concentration is varied. Other dNTPs could also inhibit the *G. intestinalis* dAK activity with a potency that mirrored the efficiencies of the corresponding deoxyribonucleosides shown in Fig. 1C and Table 1; dATP was the strongest inhibitor whereas a much weaker effect was obtained with dGTP and dCTP (Fig. 1F). As expected from this rationale, dTTP did not inhibit the reaction.

**Table 1.** Enzyme kinetics of *G. intestinalis* dAK with different substrates. The truncated dAK is described in the mutational analysis section. The values indicate the average with standard errors from three independent experiments. The k_cat_ value is shown as activity per polypeptide. Abbreviations: Ara-A, adenine arabinoside, FANA-A, 2’
s-fluoro-β-D-arabinosyl adenine (additional F- and Cl labels indicates modifications in the 2 position of the adenine moiety).

| Substrate | $K_M$ ($\mu\text{M}$) | $V_{\text{max}}$ ($\mu\text{mol}\cdot\text{min}^{-1}\cdot\text{mg}^{-1}$) | $k_{\text{cat}}$ ($\text{s}^{-1}$) | $k_{\text{cat}}/K_M$ ($\text{M}^{-1}\cdot\text{s}^{-1}$) |
| --- | --- | --- | --- | --- |
| dAdo | $1.12 \pm 0.37$ | $10.3 \pm 1.0$ | 4.96 | $4.43 \times 10^6$ |
| dGuo | $476 \pm 133$ | $4.87 \pm 0.49$ | 2.36 | $4.94 \times 10^3$ |
| dIno | $701 \pm 83$ | $9.49 \pm 1.14$ | 4.59 | $6.54 \times 10^3$ |
| dCyd | $2090 \pm 380$ | $12.1 \pm 1.3$ | 5.87 | $2.81 \times 10^3$ |
| dThd | | $<0.25^*$ | | |
| dUrd | | $<0.1^*$ | | |
| Cladribine (Cl-dAdo) | $0.51 \pm 0.08$ | $5.78 \pm 1.16$ | 2.79 | $5.48 \times 10^6$ |
| Ara-A | $6.70 \pm 0.07$ | $6.60 \pm 0.06$ | 3.19 | $4.76 \times 10^5$ |
| F-Ara-A | $8.40 \pm 0.60$ | $7.20 \pm 0.82$ | 3.48 | $4.14 \times 10^5$ |
| FANA-A | $2.66 \pm 0.11$ | $7.64 \pm 0.64$ | 3.69 | $1.39 \times 10^6$ |
| Clofarabine (Cl-FANA-A) | $6.23 \pm 1.33$ | $7.26 \pm 1.32$ | 3.51 | $5.63 \times 10^5$ |
| **Truncated dAK (30-232)** |  |  |  |  |
| dAdo | $53.8 \pm 3.10$ | $13.0 \pm 1.9$ | 5.22 | $9.69 \times 10^4$ |
| dCyd | $31600 \pm 6600$ | $11.0 \pm 1.7$ | 4.40 | 139 |

\*Numbers indicate the detection limit in activity assays with 2 mM substrate.

### *G. intestinalis* dAK has a high catalytic efficiency with deoxyadenosine and analogues modified at position 2 and 2’ s

Enzyme kinetic parameters of the *G. intestinalis* dAK activity were determined with natural deoxyribonucleosides as well as deoxyadenosine analogues (Table 1). Among the natural deoxyribonucleosides, deoxyadenosine stands out by its low K_M_ value whereas the differences in V_max_ was much less pronounced. Substrate selection is thus primarily affinity-based with deoxyadenosine being strongly preferred over the others. However, it cannot be excluded that the low-affinity substrates may be relevant as well in the cell where other factors such as transport efficiency and further metabolism into the triphosphate form can play important roles. Table 1 also shows the efficiency of different deoxyadenosine analogues as substrates. To validate the results with the deoxyadenosine analogues shown in Table 1, we also made an alternative assay where each analogue was tested at 200 μM concentration with and without 200 μM deoxyadenosine as competitor (Fig. 2). The substrate concentration was chosen to be well above the K_M_ for all substrates. The results in Fig. 2 matched well with the conclusions in Table 1. When tested alone, the analogues gave activities that were very similar to the V_max_ values and in the competition experiments, deoxyadenosine could efficiently compete out the high-K_M_ substrates (>90% inhibition) but had less than 50% effect on the 2-halogenated versions of deoxyadenosine (45% effect with cladribine and 33% with F-dAdo). F-dAdo was not included in Table 1 due to that the HPLC-based assay was not sensitive enough to obtain reliable K_M_ values below 1 μM, but for cladribine we could determine the K_M_-value via an alternative radiolabeled filter assay showing that it had the highest affinity of the substrates tested. However, the difference in K_M_ as compared to deoxyadenosine was modest and, as described above, the competition assay indicated nearly equal affinities of the two substrates. The beneficial effect of 2-halogenation is thus not comparable to the 20-fold improvement reported for the human dCK that is responsible for deoxyadenosine phosphorylation in the cytosol [25]. Nevertheless, *G. intestinalis* dAK is generally a much more active enzyme than the human dCK and the catalytic efficiency is >100-fold higher with cladribine and >1000-fold higher with deoxyadenosine. A general trend for the deoxyadenosine analogues in Table 1 is that the K_M_ value for deoxyadenosine is lower than the corresponding arabinosyl- and 2’
s-F-substituted arabinosyl derivatives (Ara-A and FANA-A), which was in line with the conclusions from the competition assay. Nevertheless, all deoxyadenosine analogues tested had catalytic efficiencies within approximately one order of magnitude and can thus be considered good substrates. The effect of 2-halogenation varied but was never as prominent as for the mammalian dCK and was inhibitory in the case of clofarabine (Table 1).

**Fig. 2.**
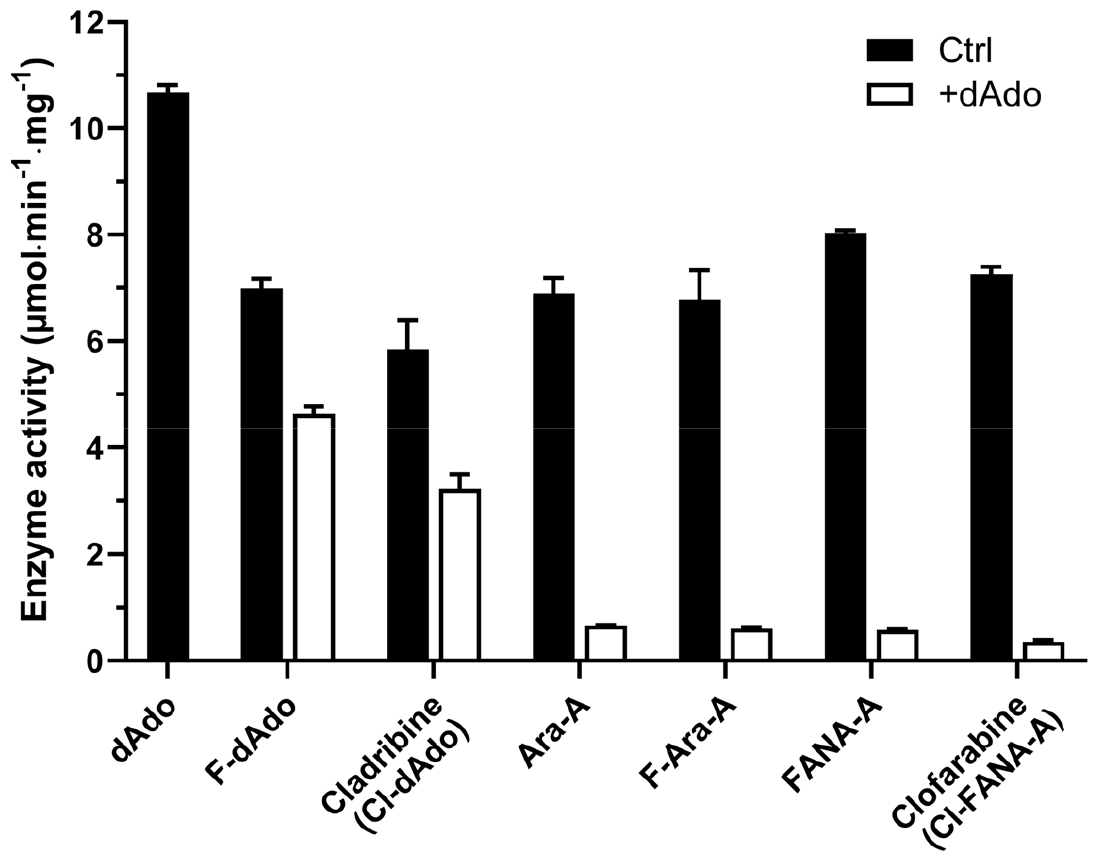
*G. intestinalis* dAK is active with deoxyadenosine analogues modified at the 2 and 2’
spositions. The activity assays were performed with 200 μM analogue in the absence or presence of 200 μM deoxyadenosine to get information about both activity and affinity. The results are based on three independent experiments with standard errors indicated.

### Deoxyadenosine analogues inhibits *G. intestinalis* proliferation and encystation

A growth inhibition assay with different deoxyadenosine analogues showed that they inhibited the *G. intestinalis* parasites with EC_50_ values that mirrored the substrate specificity of dAK. The EC_50_ values of the analogues on the WB strain were in the range from 1-7 μM (Table 2), and comparable to metronidazole which gives an EC_50_ value of 2.7 μM on the same strain under these conditions [19]. Among the analogues tested here, cladribine was the most active one, whereas a higher selectivity as compared to the mammalian reference cell line was obtained with either Ara-A or F-Ara-A (Table 2 and S4 Fig). A main contributor to the selectivity of Ara-A is that the drug is deaminated by the mammalian cells but not the parasites. Indeed, our previous analysis shows an EC_50_ value of 1.8 μM on Balb/3T3 fibroblasts (and >10 μM on human WS1 fibroblasts) when the drug is combined with 1 μM EHNA to inhibit mammalian adenosine deaminase [26]. In the case of cladribine and F-Ara-A, they are deamination resistant and results with these analogues are not dependent on the presence of adenosine deaminase inhibitors. All analogues tested were fully active on metronidazole-resistant *G. intestinalis* strains as well, which is important for their potential as drugs against giardiasis. Cladribine was also verified to nearly completely inhibit the ability of the parasites to form cysts if added in the beginning of the encystation process and reduced the number of mature cysts with a 16N genome if added later (S5-S6 Fig). A lower number of 16N cells is indicative of inhibited DNA replication during encystation. In conclusion, the experiments on cells provide a proof of principle that deoxyadenosine analogues can inhibit the pathogen in the low micromolar range and are efficient regardless of cellular metronidazole-resistance.

**Table 2.**
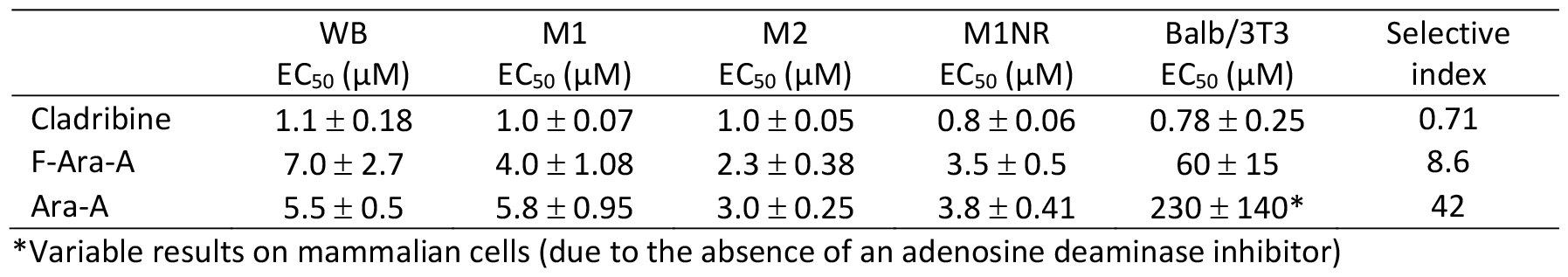
EC_50_ values showing the inhibition of *G. intestinalis* and mammalian fibroblast (Balb/3T3) proliferation. The experiments were performed with wild-type *G. intestinalis* cells (WB strain), metronidazole-resistant cells (M1 and M2) and resistance-revertant cells (M1NR). The values represent the results from three independent experiments with standard errors indicated.

|  | WB<br>EC<sub>50</sub> (μM) | M1<br>EC<sub>50</sub> (μM) | M2<br>EC<sub>50</sub> (μM) | M1NR<br>EC<sub>50</sub> (μM) | Balb/3T3<br>EC<sub>50</sub> (μM) | Selective<br>index |
| --- | --- | --- | --- | --- | --- | --- |
| Cladribine | 1.1 ± 0.18 | 1.0 ± 0.07 | 1.0 ± 0.05 | 0.8 ± 0.06 | 0.78 ± 0.25 | 0.71 |
| F-Ara-A | 7.0 ± 2.7 | 4.0 ± 1.08 | 2.3 ± 0.38 | 3.5 ± 0.5 | 60 ± 15 | 8.6 |
| Ara-A | 5.5 ± 0.5 | 5.8 ± 0.95 | 3.0 ± 0.25 | 3.8 ± 0.41 | 230 ± 140* | 42 |

\*Variable results on mammalian cells (due to the absence of an adenosine deaminase inhibitor)

### The *G. intestinalis* dAK structure reveals a novel dimerization interface

To structurally characterize the active site and investigate the substrate preference of *G. intestinalis* dAK, we solved the structure of full-length *G. intestinalis* dAK by macromolecular X-ray crystallography (Fig 3A, S1 Table). The asymmetric unit (ASU) contained two copies of *G. intestinalis* dAK, of which each monomer showed a similar fold to the homologous structure of *Mycobacterium mycoides* dAK (*M. mycoides* dAK), as well as human dGK and dCK (S7 Fig) [27-29]. However, when superimposed, the structure revealed an alternative, additional dimer assembly than formerly seen for *M. mycoides* dAK (Fig. 3B). This novel interface suggested that *G. intestinalis* dAK could be a tetramer, which is stabilized by the extended N- and C-termini of the monomers (Fig 3C). Here, the N-terminal residues (M24_MI_-P29_MI_) of the first monomer (MI) form extensive interactions with residues of the second monomer (MII) (Y112_MII_, R119_MII_, Y132_MII_, F192_MII_, R199_MII_) (Fig 3D, S2 Table). Hydrogen bonds are formed between the sidechains and amide backbone oxygen of Y132_MI_:M24_MI_, R119_MII_:M24_MI_, Y132_MII_:G25_MI_, and R199_MII_:P29_MI_ whereas Y112_MII_ and F192_MII_ together with F26_MI_, P27_MI_, and Y28_MI_ form a hydrophobic pocket (Fig 3D). The C-terminus of MI forms an intermolecular β-sheet with the twisted four-stranded β-sheet of MII in addition to a salt bridge between D232_MI_ and R204_MII_, and H-bonds between D156_MII_:T240_MI_, Y186_MII_:T240_MI_, A208_MII_:D236_MI_, Y210_MII_:D236_MI_, R211_MII_:D236_MI_, and H227_MII_:H227_MI_ (Fig 3D). Thus, the *G. intestinalis dAK* structure reveals a novel, hitherto unseen, dimerization interface.

**Fig 3.**
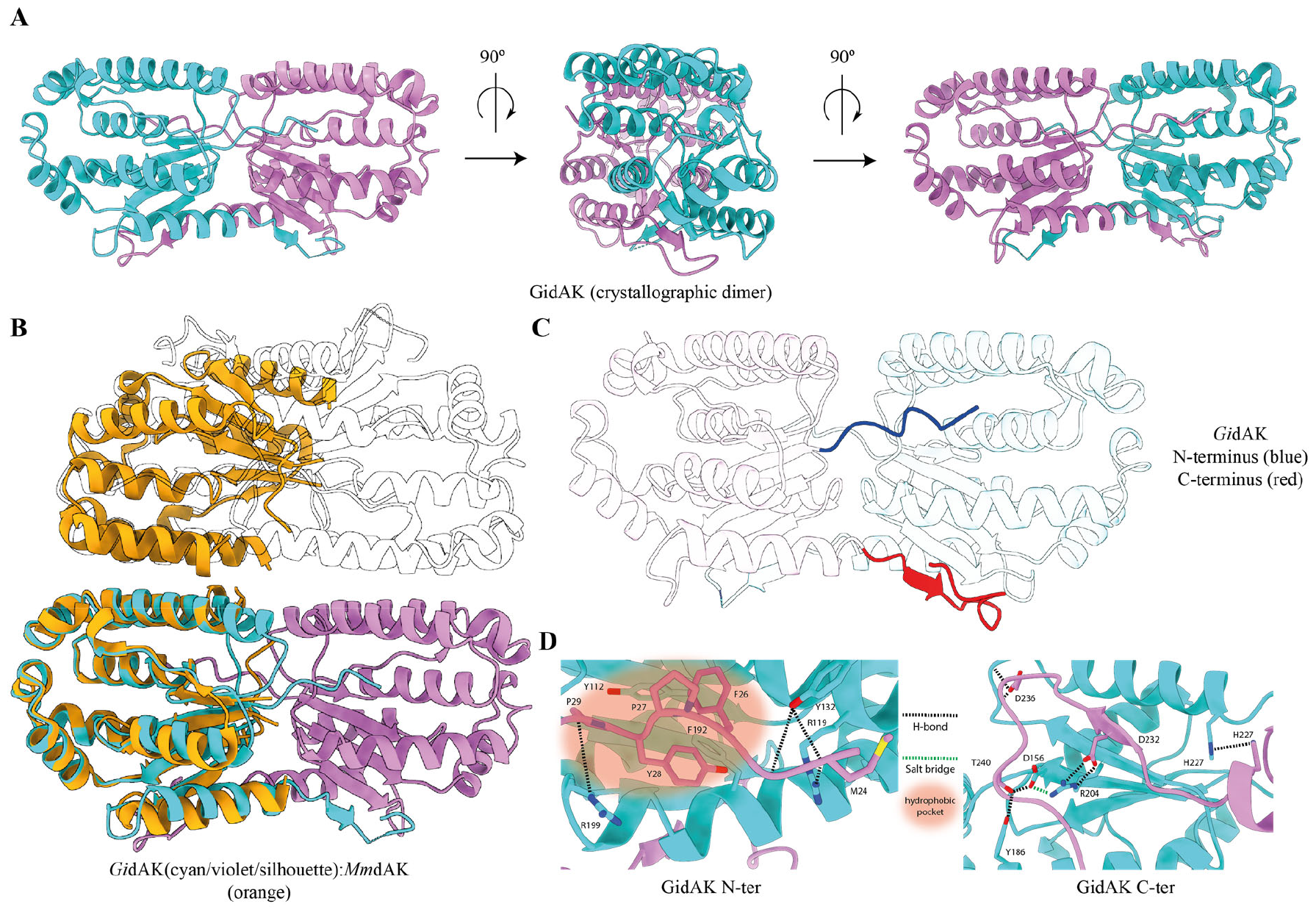
The crystal structure of *G. intestinalis* dAK reveals a novel dimerization interface. (A) Three different views of the crystallographic dimer of *G. intestinalis* dAK rotated horizontally counterclockwise by 90°. (B) The crystallographic dimer of *G. intestinalis* (cyan/violet) superimposed on the structure of *M. mycoides* dAK (orange, PDBID: 2JAQ). The crystallographic dimer exhibits a novel interface whereas the symmetry-related dimer (shown as a silhouette) is oriented in a similar way as in the *M. mycoides* dimer. This indicates a tetrameric assembly of the *G. intestinalis* dAK (cyan/violet/silhouette). (C) Illustration showing the key roles of the N-terminus (blue) and C-terminus (red) in the crystallographic dimer assembly. (D) Detailed representation of the of N-(left) and C-termini (right) interaction pattern with the neighboring monomer to allow for the dimerization. N-terminal interactions: hydrophobic pocket: F26_MI_, Y28_MI_, P27_MI_, P29_MI_, Y112_MII_, F192_MII_; hydrogen bonds: M24_MI_:R119_MII_, Y132_MII_, G25_MI_:Y132_MII_, P29_MI_:R199_MII_. C-terminal interactions: Salt bridges: D232_MI_:R204_MII_; hydrogen bonds: H227_MI_:H227_MII_, D236_MI_:R206_MII_, A208_MII_, Y210_MII_, T240_MI_:D156_MII_, Y186_MII_. (MI: monomer 1, MII, monomer 2). The hydrophobic pocket is shown as a transparent orange oval, hydrogen bonds as black dotted lines, and salt bridges as green dotted lines. Protein chains are shown in cartoon representations and interacting residues as sticks.

### Co-purified deoxyadenosine mono- and diphosphate ligands bound to *G. intestinalis* dAK allow for the modelling of substrate analogues

After structure solution by molecular replacement, we identified unbiased *F*_*o*_*-F*_*c*_ density that corresponds to ligands bound in the active sites of the crystallographic dimer. Interestingly, no substrates or analogues had been added during protein purification or during crystallization, suggesting that the bound ligands were present already in the cell extract and exhibit a high binding affinity for dAK. The ligand density was well enough resolved to identify shapes corresponding to an adenine and a deoxyribose sugar. However, the number of phosphate groups differed in the two monomers: whereas a dADP, with a partial occupancy of 0.5 for the β-phosphate, could be modelled into active site I, the density for active site II unambiguously allowed for the placement of a dAMP (S8 Fig). HPLC analysis of the dAK preparation showed that the protein contains 0.40 dAMP and 0.35 dATP per polypeptide, and subsequent analysis showed that dATP could slowly be converted to dADP by the enzyme preparation over a matter of days at room temperature (S9 Fig). The dATP-dephosphorylating activity was several orders of magnitude lower than the regular dAK activity and could possibly be the result of other enzymes present as minor impurities in the protein preparation. The active site residues of *G. intestinalis* dAK are highly conserved to the residues found in *M. mycoides* dAK (S1 Fig), with the adenine base being held in place by face-to-edge (F72) and face-to-face (F117) π-stacking, and hydrogen bonds between the adenine nitrogen N1, N3, and N10 and Q84, an H_2_O molecule, and Q84 and D114, respectively. The deoxyribose sugar oxygen forms hydrogen bonds with an H_2_O molecule and the 3’-hydroxy group forms hydrogen bonds with E171 and Y73. Finally, the α-phosphate forms hydrogen bonds with G40, K43, R109 via an H_2_O molecule, and R169 (Fig 4A). Our biochemical studies indicated that cladribine is a slightly better substrate than deoxyadenosine. By using the adenine base as reference, we therefore modelled cladribine as well as all other tested substrate analogues into the active site of *G. intestinalis dAK* (Fig 4B). The modelled cladribine ligand suggested that the 2-Cl group points towards F72, altering the face-to-edge π-stacking to a halogen-π interaction (Fig 4B), which has been shown to be a stronger type of interaction [30, 31]. This provides a possible explanation for how cladribine and F-dAdo can have a higher affinity than deoxyadenosine. However, the model did not demonstrate a perfectly perpendicular face to edge interaction and the gain in affinity compared to deoxyadenosine was modest (Table 1 and Fig. 2), indicating a potential for ligand improvement in drug development.

**Fig 4.**
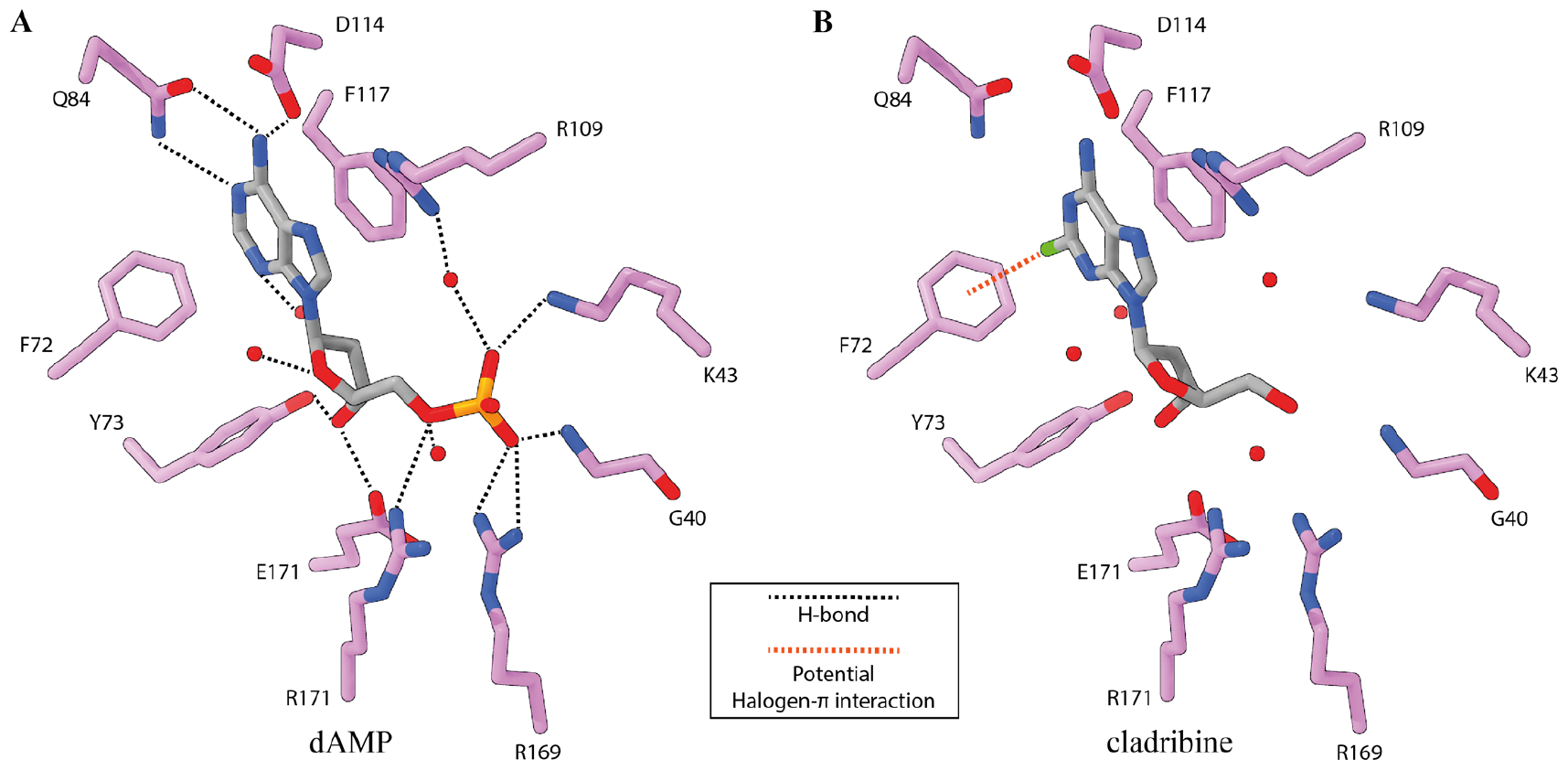
The active site of *G. intestinalis* dAK reveals an extensive hydrogen bond pattern. (A) Stick representation of dAMP (grey sticks) bound by active site residues (magenta sticks) of *G. intestinalis* dAK. Water molecules are shown as red spheres and hydrogen bonds as black dotted lines. (B) Representation of cladribine (grey sticks) modelled into the active site of *G. intestinalis* dAK, using the adenine base as reference. A potential, additional halogen…π-interaction is shown as an orange dotted line.

### Cryo-electron microscopy reveals that dAK exists as a tetramer in solution

The crystal structure suggested that dAK may be present as a homotetramer in solution (Fig. 5A). To ascertain whether the interactions in the putative homotetramer were due to the crystal packing or reflecting the true state of the enzyme in solution, we used cryo-electron microscopy (cryo-EM) single particle analysis (S10 Fig). The experimental 2D class averages obtained by cryo-EM closely resembled computer-generated 2D re-projections of the putative tetramer from the crystal structure (Fig 5B). Due to strong preferred orientations, it was not possible to get a high-resolution 3D structure of dAK using cryo-EM and the resulting 3D map therefore had an anisotropic appearance (S11 Fig.). However, the 3D map was sufficient to perform an approximate fitting of the crystal structure tetramer, whose size and shape fitted the cryo-EM map well at the attained resolution (Fig. 5C). Taken together, cryo-EM thus clearly confirmed that dAK exists in solution as the tetramer suggested by the x-ray crystallography data.

**Fig 5.**
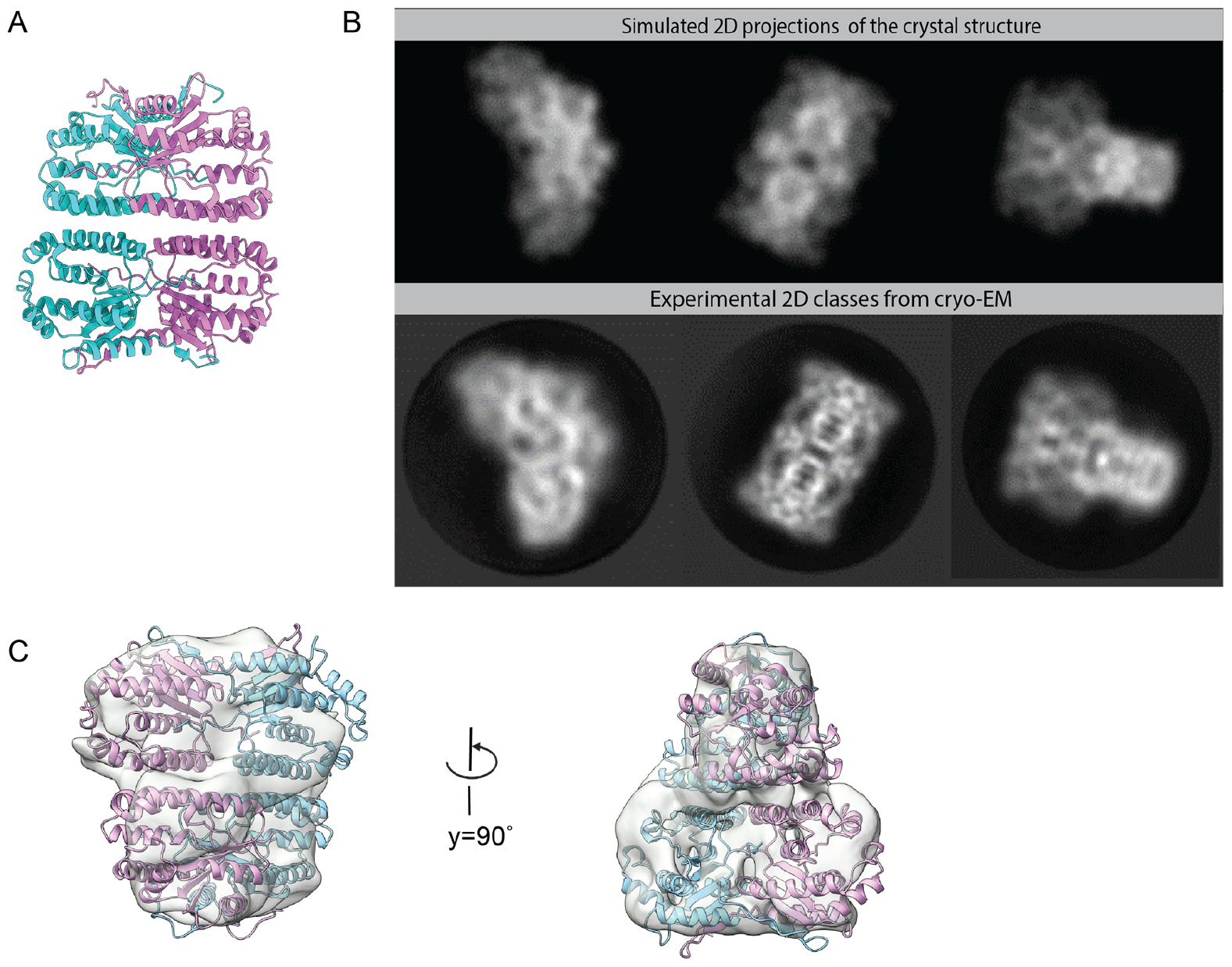
Cryo-EM shows that *G. intestinalis* dAK exists as a tetramer in solution. (A) Crystal structure of *G. intestinalis* dAK, showing the suggested homotetramer quaternary structure. The canonical homodimers found in previous structures from the non-TK1-like family are colored in cyan and violet. (B) The top row shows computational simulations (reprojections at 7 Å resolution) of the 2D class averages that the crystal structure homotetramer would lead to. The lower row shows corresponding experimental 2D class averages from the cryo-EM data. (C) The crystal structure as shown in panel A, fitted in the anisotropic 3D density map obtained by cryo-EM (C1 symmetry, nominal resolution 4.8 Å).

### Deletion mutants of *G. intestinalis* dAK show that the N- and C-termini are important for tetramerization and substrate affinity

In order to study the functional relevance of tetramerization, a deletion mutant of dAK was created that lacked residues 1-29 and 233-251 of the N- and C-termini (dAK-ΔNΔC), which were indicated to be important for subunit interaction in the crystal structure. Size exclusion chromatography indicated that the deletion mutant indeed formed a structure considerably smaller than a tetramer. However, the estimated molecular mass of 71 kDa is in between a dimer (45 kDa) and a tetramer (90 kDa) based on comparison with standard proteins (Fig. 6A). The peak exhibited considerable tailing indicating that the mass estimation was not reliable and could be the effect of equilibration between different forms or interaction with the chromatography material. In contrast, the wild-type protein gave a symmetrical peak and clearly indicated a tetramer (theoretically 116 kDa). Using mass photometry, it was possible to get clearer results with the deletion mutant. This method requires much lower protein concentrations (typically <100 nM) than size exclusion chromatography. The wild-type protein still behaved as a tetramer at 65 nM concentration, whereas measurements of the deletion mutant clearly indicated that it is predominantly a dimer at this protein concentration (Fig. 6B). The results supported the view that the size exclusion chromatography (run at 4 μM protein) may result in a dimer-tetramer equilibrium that almost completely shifts to dimer at the lower protein concentration used in mass photometry (65 nM). Mass photometry experiments on other deletion mutants lacking either the N- or C-terminus showed that they predominantly formed tetramers but with a weaker structure as indicated by a significant proportion of dimers (Fig. 6B). In contrast, the dimers were undetectable in wild-type dAK samples. Enzyme activity assays with the double deletion mutant (dAK-ΔNΔC) showed that the lost ability to readily form tetramers was accompanied by a ∼50 times reduced affinity for deoxyadenosine (i.e. higher K_M_) but no major change in the k_cat_ value (Table 1). The results indicate that it is only the substrate affinity and not the catalytic mechanism that is dependent on the ability to form tetramers. Corresponding experiments with deoxycytidine as an example of a structurally different substrate showed a similar trend with the K_M_ value being much more affected than k_cat_. Thus the reduced deoxyadenosine affinity in the double-deletion mutant seems to be a general loss of substrate affinity rather than a change in specificity.

**Fig 6.**
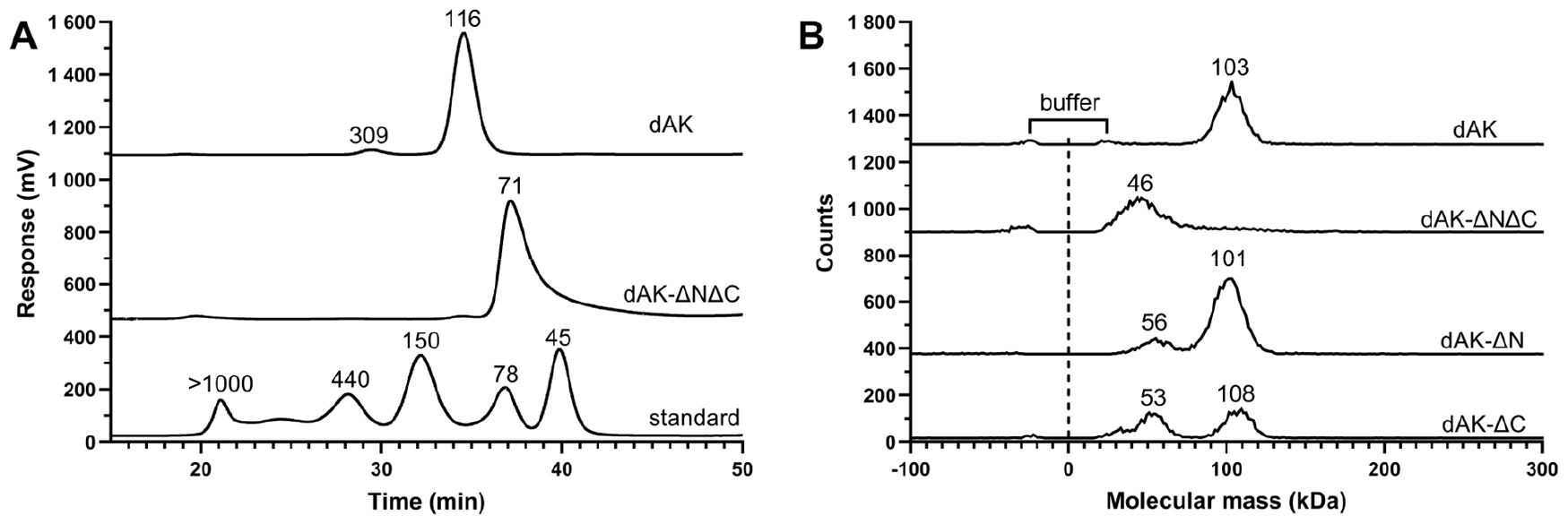
Size exclusion chromatography and mass photometry analysis of full-length and truncated *G. intestinalis* dAK confirm that the N- and C-termini are important for tetramer formation. (A) Size exclusion chromatography of full-length dAK, truncated dAK lacking the N- and C termini (dAK-ΔNΔC) and a protein standard. (B) Mass photometry experiments of wild-type dAK, dAK-ΔNΔC, dAK-ΔN and dAK-ΔC. The molecular masses of the dAK peaks in A-B were assessed by comparing them to a standard curve of proteins with known molecular masses. In B, the mass assessment was preceded by first fitting the peaks to Gaussian curves by the instrument software. The buffer peak indicated in B deviates from a typical protein peak by having equally sized positive and negative peaks coming from binding and unbinding events, respectively. The different graphs in A-B are shifted along the ordinate in order to fit many experiments in the same panel.

## Discussion

The ability to live without ribonucleotide reductase seems to be a daunting task considering how few organisms have succeeded. There are several challenges to overcome, with the most obvious one being the scarce supply of substrates for the salvage synthesis of dNTPs. The concentration of deoxyribonucleosides is generally below the detection limit in most environments and none of the ribonucleotide reductase-lacking organisms found so far is free-living. In essence, it should be possible to degrade DNA from dead cells in the surroundings as a source for salvage but given that no free-living organism has succeeded to live entirely on salvage, it is likely to be difficult and it should be remembered that also among pathogens it is very few that have succeeded.

*G. intestinalis* seems to be adapted for the problem of scarce sources or deoxyribonucleosides in several ways. The location of the parasites in the duodenum is one important factor. In this part of the gastrointestinal system, the food is processed and thereby available for the parasite and there are also few other organisms that it needs to compete with. Nevertheless, it is still a challenge to compete with the mammalian cells for the absorption of the deoxyribonucleosides and it therefore needs an efficient system for uptake and salvage. In the case of *G. intestinalis* dAK, this has been accomplished by having a >1000 times higher catalytic efficiency (k_cat_/K_M_) than the human dCK and deoxyguanosine kinase (dGK) have for deoxyadenosine [25, 32, 33]. The latter enzyme is responsible for the phosphorylation of purine deoxyribonucleosides in the mitochondria [17]. In fact, none of the studied dNKs up to date has such a high catalytic efficiency for deoxyadenosine as the enzyme from *G. intestinalis*. The high substrate affinity of *G. intestinalis* dAK goes hand in hand with the previous studies of the thymidine kinase of the parasite, which has a much higher affinity for thymidine than mammalian TK1 (K_M_ values: 0.07 vs 0.5 μM) [19]. It is typical that members from the TK1-family have higher affinities than non-TK1-like dNKs, and the K_M_ of the *G. intestinalis* thymidine kinase needs therefore to have an exceptionally low K_M_ to be able to compete with the mammalian enzyme. The current study indicates that it could be a general trend that *G. intestinalis* has developed high-affinity dNKs to be able to compete with its host although it remains to be discovered which enzyme is responsible for the phosphorylation of deoxycytidine and deoxyguanosine in the parasite. Although dAK has comparable k_cat_ values for these substrates as for its main substrate, the affinity for them is much lower and it is therefore more likely that the second, yet not studied, non-TK1-like dNK in the parasite is responsible. In conclusion, the parasite seems to have developed highly competitive enzymes for deoxyribonucleosides, at least for deoxyadenosine and thymidine. In the case of deoxyadenosine, a K_M_ of 1.1 μM is low enough to be competitive in comparison to the corresponding mammalian enzymes, whereas for thymidine a much lower K_M_ is required (0.07 μM).

A second challenge for *G. intestinalis* is how to regulate the salvage of deoxyribonucleosides. A hint of how important that is comes from a study of an engineered *E. coli* cell line lacking all ribonucleotide reductase genes and supplemented with the *M. mycoides* dAK gene to complement its own salvage and be able to proliferate in the presence of deoxyribonucleosides in the medium [34]. Over time the cell line downregulated deoxyribonucleotide degradation pathways and gradually evolved the ability to live on lower and lower concentrations of deoxyribonucleosides in the medium. However, it came with the cost of a substantially increased mutation rate. This highlights the key role of ribonucleotide reductase not only to produce deoxyribonucleotides but also for the regulation of the process to keep the dNTP pools balanced and minimize replication errors. It is an enigma how *G. intestinalis* is able to balance the different dNTP pools, but one part of the explanation could be its strong feedback inhibition by dATP (K_i_= 0.34 μM). This feedback inhibition will efficiently turn the enzyme activity off if dATP is not immediately used up for DNA synthesis. Another advantage of the strong dATP inhibition is to secure that the substrate is spread equally among the cells in the population. With uncontrolled salvage, there is a risk that the first parasites that encounter the substrate will take the major share and leave too little for the others.

The finding of the much higher catalytic efficiency of *G. intestinalis* dAK for deoxyadenosine as compared to the mammalian dNKs is valuable for drug discovery. One way to target the dependency on salvage would be to make inhibitors against dAK, but that requires development of drugs from scratch, and we have instead focused on substrate analogues which after phosphorylation by the enzyme target downstream processes. The potential advantage of using substrate analogues is then both based on the higher substrate affinity of *G. intestinalis* dAK as compared to the mammalian dNKs and on the fact that the parasite cannot become resistant by downregulating dAK without also losing its ability to replicate. Clinically used deoxyadenosine analogues are primarily targeting cancer cells or virus-infected cells. The benefit of using already known drugs is that many of the pharmacological properties are already known and fewer clinical trials are needed before it can become a final drug against the parasite. The three deoxyadenosine analogues most commonly used against cancer are cladribine, fludarabine and clofarabine, which block cell division by inhibiting ribonucleotide reductase and interfering with DNA replication. They are all halogenated at position 2 of the nucleobase to make them stable against deamination by the mammalian adenosine deaminase. The analogues can be given orally as well as intravenously with the latter option giving higher blood concentrations for a given dose, which is generally preferred in anticancer treatment. In the case of *G. intestinalis*, oral treatment is more relevant due to the localization of the parasite in the duodenum. An advantage is then that the drugs will not be diluted in the blood before they can reach the parasites, which hopefully means that lower doses can be used.

*G. intestinalis* differ from the mammalian cells in several ways of importance for the development of deoxyadenosine analogues. First, the parasite lacks adenosine deaminase and it is therefore less critical that the drug is protected against deamination although some protection may still be needed for the drug to be stable enough to reach the site of infection. Second, *G. intestinalis* also lacks ribonucleotide reductase and it is therefore only the direct interference on DNA replication that is relevant for the antigiardial effect. Our experiments demonstrated that the parasite was sensitive to deoxyadenosine analogues that effectively inhibited proliferation as well as encystation. Cladribine (Cl-dAdo) had the highest catalytic efficiency with *G. intestinalis* dAK of the drugs tested, and modeling of the substrate analogue into the *G. intestinalis* dAK structure indicated that a halogen…π interaction between Phe72 in the protein and the chlorine at position 2 in the substrate contributed to the affinity (Fig. 4B). However, the 2-chlorination only gives a 2-fold increased affinity as compared deoxyadenosine and virtually no change in catalytic efficiency (the k_cat_ is lower). The effect is thus modest as compared to the mammalian dCK and dGK where 2-chlorination leads to a 10-25-fold increased catalytic efficiency [25, 33]. Although this is far from compensating the superior catalytic efficiency of *G. intestinalis* dAK, it should be remembered that the parasites lack one of the drug targets (ribonucleotide reductase) and experiments on cells showed nearly identical EC_50_ values on *G. intestinalis* as compared to mammalian fibroblasts. Better selectivity could instead be obtained with the two drugs F-Ara-A and Ara-A. Similarly to cladribine, F-Ara-A is 2-halogenated but with the advantage of not being as good substrate for mammalian dCK and dGK [33, 35]. Ara-A, which is an even worse substrate of dCK and dGK and is mainly phosphorylated by adenosine kinase in mammalian cells, could perhaps be the most interesting option of the two drugs against giardiasis. Clinically it has been used as a topically applied drug against herpes simplex virus and is generally regarded non-toxic if ingested due to the rapid deamination in the body (https://www.rxlist.com). Ara-A (as well as F-Ara-A) is generally given in a phosphorylated form (Ara-AMP) to increase its solubility and needs to be dephosphorylated in the body before it can be taken up by cells. An advantage for the treatment of giardiasis is that the phosphorylated form of the drug is protected against adenosine deaminase, which increases the chances that it can reach the site of the parasites intact. Once there, Ara-A will be liberated from Ara-AMP by intestinal phosphatases. If needed for stability, the drug can also be combined with an adenosine deaminase inhibitor. However, then it is important to be aware that conventional adenosine deaminase inhibitors used as anticancer drugs are optimized for a strong systemic inhibition of the enzyme and associated with side effects. In the case of giardiasis, it is sufficient to inhibit the enzyme in the local intestinal environment for a limited time and then weaker inhibitors can be considered. A wide range of non-toxic substances that inhibits the enzyme has been described, including garlic and other plant extracts [36].

The crystal structure of *G. intestinalis* dAK sheds light into how the enzyme has attained such a high affinity to its main substrate and how it can be exploited for drug discovery. From a drug development viewpoint, more selective nucleoside analogues can be developed by taking advantage of the enzyme structure. The mammalian dCK does for example not use a halogen-π interaction with the 2-chloro atom in clofarabine and the difference between the enzymes means that it may be possible to develop analogues with other 2-modifications that fit better with the *G. intestinalis* dAK than the mammalian enzyme. Especially, it would be valuable to avoid the beneficial effect that the 2-halogen has on binding to the mammalian dCK and dGK.

The structure of the *G. intestinalis* dAK also gave insights into how such a strong affinity for deoxyadenosine has been obtained. This is the first example of a non-TK1-like dNK with a tetramer structure and subsequent mutagenesis analysis showed that the tetramer structure and the affinity are connected. By deleting the N- and C-termini and thereby breaking up the enzyme into dimers, the enzyme lost its high substrate affinity. The V_max_ was not changed significantly indicating that the enzyme did not lose its ability to catalyze the reaction. Interestingly, the enzyme lost affinity for both deoxyadenosine and deoxycytidine indicating that this was a general effect and not a change in specificity. The ability to gain substrate affinity by tetramerization could possibly represent an evolutionary shortcut explaining how the ancestor to the parasite became independent of *de novo* dNTP synthesis. For comparison, changes more directly associated to the active site are likely to cause undesired side effects on catalysis or specificity, which must be handled with compensatory mutations. The tetramerization is an elegant way to directly increase the affinity without affecting other parameters and may be a general feature in the *G. intestinalis* dNKs that can explain how the parasite became able to survive solely on salvage for dNTP synthesis.

## Materials and Methods

### Cloning of the *G. intestinalis* dAK into an expression vector

The identified *G. intestinalis* dAK gene in the assemblage A isolate WB (GL50803-17451 at https://giardiadb.org/giardiadb) was amplified with PCR using extracted genomic DNA from *G. intestinalis* WB as template. The PCR reaction was performed with the primers GT**C CAT GG**C TGC GCG AAA TCT GG (forward primer) and CAT GCT **GGT ACC** TTA ATA GAA ATT GGA AGC TGA TTG CG Reverse primer) with included restriction sites marked in boldface for NcoI and Acc65I, respectively. The PCR-amplified gene was cloned into the pETZ vector from European Molecular Biology Laboratory for expression in *Escherichia coli*. The resulting plasmid construct (pETZ-dAK) encoded an N-terminal His_6_-tagged fusion partner Z (10 kDa IgG-binding domain protein) followed by a tobacco etch virus cleavage site connected to the 29 kDa dAK. The obtained clones were verified to be correct by sequencing. Similar experiments were also performed to make truncated versions of the protein: dAK-ΔN (amino acids 30-251), dAK-ΔC (amino acids 1-232) and dAK-ΔNΔC (amino acids 30-232). The forward primer was then substituted by GTG **CCA TGG** CAA AGA TGT TCA TCT CTA TAT CAG G for the removal of the N-terminus and the reverse primer was substituted by GTC AGA **GGT ACC** TTA GTC TAC GAT GGC TTG GTG TTT C for the removal of the C-terminus. Both primers were substituted for the construction of the mutant lacking both termini.

### Protein expression and purification

The pETZ-dAK vector was introduced into the *E. coli* strain BL21(DE3)plys for recombinant expression of the *G. intestinalis* dAK. The bacteria were cultivated in LB medium supplemented with 50 μg/ml kanamycin and 34 μg/ml chloramphenicol at 37°C. Protein expression was induced at OD_600_ ≃0.6 with 0.5 mM isopropyl β-D-thiogalactopyranoside for overnight induction at 18°C. The cells were harvested by centrifugation at 4000 ×*g* for 15 minutes. The pellet was washed with 20 mM Tris-HCl pH 7.5, and re-centrifuged at 4000 ×*g* for 10 minutes. The supernatant was discarded, and the pellet was resuspended in lysis buffer (20 mM Tris-HCl pH 7.5, 0.4 M NaCl and 0.1 mM phenylmethylsulfonide fluoride) using a volume of 10 ml per g of pellet. The suspension was frozen and thawed, sonicated and centrifuged at 40 000 rpm for 45 minutes (Beckman L-90 ultracentrifuge using a Ti70 rotor). The supernatant was loaded onto a 2 ml column containing Ni-NTA His·Bind resin (Merck). The column was first washed with a solution containing 20 mM Tris-HCl pH 7.5, 0.4 M NaCl and 20 mM imidazole. The second wash was with 50 mM imidazole (other buffer constituents were the same). Finally, the Z-tagged dAK was eluted with a solution containing 20 mM Tris-HCl pH 7.5, 150 mM NaCl and 250 mM imidazole. In order to remove the fusion partner, the protein was first exchanged into an imidazole-free buffer containing 20 mM Tris-HCl pH 7.5 and 150 mM NaCl. TEV protease was added to the Z-tagged dAK using a ratio of 1:10, and the proteins were incubated for 3 hours at room temperature. The mixture was then loaded onto a 0.4 ml Ni-NTA His·Bind resin to bind the His_6_-tagged fusion partner and collect the *G. intestinalis* dAK in the flow-through. The protein was frozen in liquid nitrogen and stored at -80°C.

### Enzyme assay of dAK

*G. intestinalis* dAK was incubated in 50 μl of buffer containing 2 mM ATP, 5 mM MgCl_2_, 0.5 mM DTT, 100 mM potassium acetate and 50 mM Tris-HCl pH 7.5, with various concentrations of [2,8-^3^H]-deoxyadenosine at 37°C for 30 minutes. These standard conditions were based on optimization experiments (Fig. 1 and S2). After completion of the assay, the enzyme was inactivated by an incubation at 100°C for 2 minutes to stop the reaction. In this radioactive assay, the product was separated from the substrate on filters as described before [19] (modified version of previous protocol [37]). Briefly, 20 μl of each reaction was spotted onto a DE-81 filter (Whatman), dried, washed three times for 5 min with 1 mM ammonium formate and finally incubated in 1 ml HCL/KCL (0,1 M of each) with shaking for 2 min before scintillation counting. For other substrates than deoxyadenosine, the product was separated from the substrate with HPLC analysis as described below.

### HPLC analysis

Nucleotides from enzyme assays (as well as the protein preparations in S9 Fig) were analyzed using a 150 mm × 4.6 mm SunShell C18-WP HPLC column from ChromaNik Technologies Inc (Osaka, Japan) using a modified version of a previous published protocol developed for a multisubstrate assay with all five substrates (dCyd, dUrd, dThd, dGuo and dAdo) [19]. Instead of using a ternary mixture of three aqueous solutions, we changed it to a binary mixture of A and B, with the third component, tetrabuytylammonium bromide (TBA-Br) mixed into the two other solutions instead of using it separately. Solution A contained 0.7 g/l TBA-Br, 7% (v/v) methanol and 23 g/l KH_2_PO_4_, adjusted to pH 5.6 with KOH. Solution B contained the same concentration of TBA and methanol as solution A but lacked the phosphate component (B=0.7 g/l TBA-Br in 7% methanol). The samples were diluted in water and mixed with a 10× Loading buffer prior to analysis as described below, and the HPLC was run isocratically at 1 ml/min with 11% buffer A for the separation of the dNMPs from the corresponding deoxyribonucleosides. The retention times of the dNMPs were determined with a standard sample prior to setting up the HPLC program to determine a timepoint of the last peak (dAMP at ∼20 min) and based on that add an elution step coming 2 min after dAMP elution. The percentage of solution A is then quickly raised to 90% for an isocratic elution of ATP over 10 min, before quickly returning to initial conditions and equilibrated for 10 min before loading the next sample. The peaks were analyzed by UV detection at 270 nm and quantified by comparing the peak heights with a known standard. For testing deoxyadenosine and deoxyadenosine analogues in the enzyme assay, solution A and B were modified by using 5.8% (v/v) acetonitrile instead of methanol (i. e. A=5.8% acetonitrile, 0.7 g/l TBA-Br and 23 g/l KH_2_PO_4_ adjusted to pH 5.6; B=5.8% acetonitrile and 0.7 g/l TBA-Br). We then used a detection wavelength of 260 nm. Several variants of the HPLC protocol with different percentage of solution A were developed based on previously established principles for nucleotide analysis on the column [38] to be able to separate the products from ATP, ADP, the corresponding substrate and other peaks present in the sample (S3 Table). Common to all protocols was that the elution of the peaks of interest was performed isocratically, but in some cases we needed an additional elution step with 90% A as described above. Briefly, the HPLC conditions can be categorized into two groups:

- 75% A (and variants of it): for dAK assays with deoxyadenosine, F-Ara-A, F-dAdo, cladribine and clofarabine. This was suitable for assays with 200 μM substrate or more (when the samples could be highly diluted) but needed individual optimizations for the different substrates with less diluted samples as described in S3 Table.
- Low A content (7% or 20%): suitable for dAK assays with Ara-A, dAdo, FANA-A and F-Ara-A. These protocols require a subsequent switch to 90% A as described above to remove ATP and other late-eluting compounds.

As mentioned above, the samples were generally mixed with water (depending on the dilution factor desired) and 10× loading buffer (one tenth of the final volume). However, the phosphate content in the 10× loading buffer was varied depending on the percentage of solution A in the mobile phase. Generally, 1 ml buffer 10× loading buffer was prepared by mixing X μl 115 mg/ml KH_2_PO_4_ (adjusted to pH 5.6 with KOH), 500-X μl water and 500 μl 14 mg/ml TBA-Br. The value of X is then adjusted to give a final concentration of KH_2_PO_4_ that is below the corresponding concentration in the mobile phase, whereas the final TBA-Br concentration becomes the same as in the mobile phase. The X values used were the following: X=100 (protocol with 7% A), X=200 (protocol with 20% A) and X=500 (remaining protocols). The loading buffer did not contain any organic solvent and was usable for both the methanol- and acetonitrile-containing mobile phases.

### Determination of dAMP, dADP and dATP present in the dAK preparation

The procedure was similar as described previously for the analysis of nucleotides from cell extracts [38]. Briefly, 200 μl 1.15 mg/ml dAK protein was precipitated with 200 μl 10% (w/v) trichloroacetic acid and centrifuged for 1 min at 16 000 ×g. The supernatant was subsequently extracted with 576 μl of a 1:0.28 (v/v) mixture of Freon (1,1,2-trifluoro-2,2,1-trichloroethane) and N,N,N-trioctylamine for the removal of trichloroacetic acid. The upper phase was saved and diluted tenfold in water and 10× loading buffer before nucleotide quantification as described in the HPLC analysis section. A volume reduction from 400 μl to 388.6 μl by the removal of 5% TCA after Freon-trioctylamine extraction was taken into account in the calculations of nucleotide content per polypeptide [38].

### *G. intestinalis* cultivation and EC_50_ determinations

EC_50_ determinations were performed on the *G. intestinalis* WB cells (parent strain), as well as metronidazole-resistant strains (M1 and M2) and resistant revertant cells (M1-NR) [39]. The parasites were cultured in TY-I-S medium as described before [40], with the metronidazole-resistant cells grown in the presence of metronidazole, except for the last passage (24 h) prior to the start of the experiment. For EC_50_ determinations, the cells were distributed on microtiter plates, grown in the presence of various concentrations of drugs over 72 h, incubated with CellTiter-Glo reagent (Promega), counted on an Infinite M200 Pro (Tecan Group, Ltd) and analyzed by the GraphPad Prism 10.1.0 software as describe before [19]. The data were fitted to log Inhibitor vs Response curves, and standard errors were determined from the variation of 3-4 independent experiments.

### *G. intestinalis* cyst formation

The effect of cladribine on cyst formation is unpublished data from a previous study of the effect of azidothymidine on the encystation process [19]. The two negative controls (Control and DMSO), and the positive control (metronidazole) shown here for comparison are therefore identical with the previous publication, whereas the data on cladribine is new (S5-S6 Fig). Cells were resuspended to a density of ∼2.5 × 10^5^ cells and allowed to attach for 3 h in 3 ml Nunc cell culture tubes #156758 (Thermo Fisher Scientific, Waltham, MA, USA) before exchanging the medium into encystation medium. The tested drugs were added at a final concentration of 10× the determined EC_50_ value (11 μM cladribine) in the beginning of the process (0 h) or at certain time points during the encystation process (4, 8, 12 or 20 h) and the experiment was terminated at 28 h. The drug was solubilized in DMSO prior to use (total volume of 10 μl added to the cells). The negative control was handled in a similar manner and the tubes were opened, closed, and inverted but with nothing added (or 10 μl DMSO in the second control). At 28 h, the cells were centrifuged (800 ×*g*), washed twice with sterile H_2_O, resuspended, and incubated for 5 days with H_2_O to make the cells permeable to dyes. The cells were finally stained with fluorescein diacetate (FDA) to measure if they were alive and with propidium iodide (PI) to measure DNA content as described before [19, 41]. Completed cysts are permeable only to FDA, whereas cells caught in the middle of the encystation process are FDA negative but permeable to propidium iodide.

### Crystallography, structure solution, and analysis

A solution of 10 mg/mL *G. intestinalis* dAK in 20 mM Tris-HCl pH 7.5, 150 mM NaCl was used to set up sitting-drop vapor diffusion crystallization experiments. Crystals grew in the preformulated Morpheus [42] H3 condition (0.1 M amino acids, 0.1 M Buffer 1 pH 6.5, 30% GOL_P4K) within 1-3 weeks. Crystals were cryo-protected by short immersion in a fresh drop of crystallization condition and flash-frozen in liquid nitrogen. Data were collected at the MAXIV BioMAX beamline [43-45], processed with XDS [46] and scaled to 2.1 Å using aimless [47] within the CCP4 suite [48]. 5% of total reflections were set aside as an *R*_*free*_ test set. Crystals belonged to space group P3221 with two molecules per asymmetric unit, a solvent content of 69.3% and a Matthews coefficient of 4.00. All subsequent data processing was performed within the PHENIX suite [49]. The structure was solved with Phaser [50], using a processed AlphaFold-predicted *G. intestinalis* dAK structure [51] as a search model. The structure was fully refined, and the model built using iterative cycles of phenix.refine [52] and COOT [53], respectively. Ligand topology was generated using eLBOW [54]. Data collection and refinement statistics can be found in the supplementary material (S1 Table). Dimer-dimer interactions were predicted using the PISA server [55].

### Molecular graphics and visualization

Figures of protein structures and electron densities were generated using ChimeraX [56]. Ligand topology was generated using LigPlot+ [57].

### Cryo-EM sample preparation

3 μl of dAK (1 mg/ml) was applied to QUANTIFOIL Cu R300 1.2/1.3 (Electron Microscopy Sciences, catalog no. Q3100CR1.3) that had been glow discharged using a PELCO easiGlow device (Ted Pella Inc.) at 15 mA for 30 s. The sample was applied by transferring 3 μl of sample onto the film-covered side of the grid, blotted, and plunge-frozen in liquid ethane, using a Vitrobot plunge freezer (Thermo Fisher Scientific), with the following settings: 4°C, 100% humidity, a blot force of −5, and a blotting time of 5 s.

### Cryo-EM data collection

Data were collected at an FEI Titan Krios G2 transmission electron microscope (Thermo Fisher Scientific) operated at 300 kV. The objective aperture was 100 μm. The same microscope was used for both data collections, equipped with a Gatan BioQuantum energy filter coupled to a K2 Summit direct detector (Gatan, Inc) (data collection 1), and a Selectris energy filter coupled to a Falcon 4i direct detector (Thermo Fisher Scientific) (data collection 2), respectively. Coma-free alignment was performed with AutoCtf/Sherpa for data collection 1 and EPU for data collection 2. Data were acquired in parallel illumination mode using EPU (Thermo Fisher Scientific) software at a nominal magnification of 215,000× (specimen pixel size 0.63 Å/px for K2, and 0.58 Å/px for Falcon 4i). Exposures were recorded as dose-fractionated movies in TIFF and EER format, for data collections 1 and 2, respectively. Data collection 1 resulted in 2534 movies, whereas data collection 2 resulted in 3420 movies. Complete data collection statistics are shown in S4 Table.

### Cryo-EM data processing

The processing of the two data collections are outlined in S10 Fig. For data collection 2, movies were converted from EER to TIFF using IMOD prior to processing [58]. Both datasets were processed using RELION (RELION 3.1 [59] for Data Collection 1 and RELION 4 [60] for Data Collection 2). The Beam-induced motion was corrected using RELION’s MotionCor2 implementation [61] and the per-micrograph contrast transfer function (CTF) estimated using GCTF [62] and CTFFIND4 [63] for Data Collection 1 and 2, respectively. Particles were picked using crYOLO [64] and subjected to reference-free 2D classification. Extensive attempts were made at 3D classification in RELION, but no distinct classes with superior 3D appearance to the consensus were obtained. Therefore, we performed 3D refinement using all particles from the selected 2D classes. One class from the 3D classification, low-pass filtered to 20 Å, was used as a reference model. After refinement, a soft-edge mask was created using RELION, and the map was post-processed. Data collection 1 and 2 generated qualitatively similar results. Both volumes were strongly affected by preferred particle orientations, resulting in anisotropic resolution (shown for data collection 2 in S11 Fig.). Due to the slightly different pixel sizes and different detector properties, the data sets were not combined. Data collection 1 generated slightly sharper-looking 2D class averages, that are thus shown in Fig. 5B. On the other hand, data collection 2 generated a 3D map that appeared less deformed by anisotropic resolution, and this map was thus used in Fig. 5C, and deposited at EMDB. The resolution of the unmasked and masked maps from data collection 2 was calculated using the Gold-Standard Fourier shell correlation (FSC threshold, 0.143) to be 6.3 Å and 4.8 Å, respectively (S10 Fig). The application of the internal D2 symmetry found in the crystal structure did not significantly improve map appearance, and only the unsymmetrized C1 map was used. The map was viewed and analyzed using ChimeraX, which was also used to manually fit the crystallographic tetramer to the map [65]. To construct 2D projections of the crystal structure, the PDB of the crystal structure was first converted to mrc and low pass-filtered to 7 Å followed by using e2project3d [66] to get 2D projections.

### Size exclusion chromatography

Proteins were analyzed on a Superdex 200 10/300 GL column from GE healthcare (Chicago, IL, USA) equilibrated using a mobile phase containing 150 mM KCl and 50 mM Tris-HCl pH 7.6 run at 0.4 ml/min. The proteins were diluted with mobile phase into a concentration of 0.1 mg/ml prior to use, loaded onto the column using a 100 μl sample loop, and measured by a UV detector set at 280 nm. The molecular mass of the protein was assessed by comparing the retention time to a standard curve obtained from a mixture of ovalbumin (45 kDa), transferrin (78 kDa), IgG (150 kDa), and ferritin (440 kDa).

### Mass photometry

The oligomeric state of dAK was analyzed on a Refeyn 2MP mass photometer (Refeyn Ltd, Oxford, UK). Prior to analysis, *G. intestinalis* dAK was mixed with PBS into a final protein concentration of 50 nM. For the calibration of the instrument, NativeMark Unstained Protein Standard (Thermo Fisher Scientific) was used.

## Acknowledgements

We acknowledge the Kempe Foundations for providing funds for the mass photometer and MAXIV Laboratory for the time on Beamline BioMAX under Proposal 20210468. Research conducted at MAXIV, a Swedish national user facility, is supported by the Swedish Research council under contract 2018-07152, the Swedish Governmental Agency for Innovation Systems under contract 2018-04969, and Formas under contract 2019-02496. Cryo-EM was performed at the Umeå Centre for Electron Microscopy (UCEM), a SciLifeLab National Cryo-EM facility supported by instrumentation grants from the Knut and Alice Wallenberg Foundation and the Kempe Foundations, and the Swedish Research council through the National Microscopy Infrastructure, NMI (VR-RFI 2016–00968). The research was funded by grants from the Swedish Research council (2022-00593 to AH; 2018-05851, 2021-01145, and 2023-02664 to LAC) and the Knut and Alice Wallenberg foundation through the Wallenberg Centre for Molecular Medicine Umeå to LAC.

## Data availability

Coordinates reported in this study have been deposited with the Protein Data Bank with accession code XXXX. Electron microscopy maps and half-maps have been deposited in the Electron Microscopy Data Bank with the accession code EMD-XXXXX.

