## Supplementary tables and figures for "*Giardia intestinalis* deoxyadenosine kinase has a unique tetrameric structure that enables high substrate affinity and makes the parasite sensitive to deoxyadenosine analogues"

### Supplementary material

**S1 Table. X-ray crystallography data collection and refinement statistics of *G. intestinalis* dAK.** Statistics for the highest-resolution shell are shown in parentheses. PDBID: XXXX.

| Parameter | Result |
|---|---|
| **Wavelength (nm)** | 0.976254 |
| **Resolution range (Å)** | 49.04 - 2.1 (2.175 - 2.1) |
| **Space group** | P 32 2 1 |
| **Unit cell (a = b ≠ c, α = β ≠ γ)** | 103.13 Å, 103.13 Å, 147.109 Å, 90°, 90°, 120° |
| **Total reflections** | 1069671 (90295) |
| **Unique reflections** | 53245 (5191) |
| **Multiplicity** | 20 (20.9) |
| **Completeness (%)** | 99.72 (99.12) |
| **Mean I/sigma(I)** | 15.4 (2.6) |
| **Wilson B-factor** | 50.25 |
| **R-merge** | 0.092 (1.396) |
| **R-meas** | 0.097 (1.431) |
| **R-pim** | 0.03 (1.465) |
| **$CC_{1/2}$** | 0.999 (0.802) |
| **Reflections used in refinement** | 53236 (5194) |
| **Reflections used for $R_{free}$** | 2687 (277) |
| **$R_{work}$** | 0.2000 (0.3090) |
| **$R_{free}$** | 0.2247 (0.3476) |
| **Number of non-hydrogen atoms** | 3835 |
| **Macromolecules** | 3676 |
| **Ligands** | 48 |
| **Solvent** | 111 |
| **Protein residues** | 444 |
| **RMS (bonds, Å)** | 0.017 |
| **RMS (angles, ˚)** | 0.91 |
| **Ramachandran favored (%)** | 98.85 |
| **Ramachandran allowed (%)** | 1.15 |
| **Ramachandran outliers (%)** | 0.00 |
| **Rotamer outliers (%)** | 0.77 |
| **Clashscore** | 3.80 |
| **Average B-factor** | 54.22 |
| **Macromolecules (B-factor)** | 54.24 |
| **Ligands (B-factor)** | 48.60 |
| **Solvent (B-Factor** | 56.08 |

**S2 Table. List of interactions in the *G. intestinalis* dAK structure**. Detailed list of predicted interactions formed between the monomers within the crystallographic dimer of the *G. intestinalis* dAK crystal structure based on calculations using the PISA server.

| Interaction # | Monomer I | Atom | Distance [Å] | Monomer II | Atom | Interaction type |
|---|---|---|---|---|---|---|
| **1** | M24 | O | 2.83 | R119 | NH1 | Hydrogen bond |
| **2** | M24 | O | 3.76 | Y132 | OH | Hydrogen bond |
| **3** | G25 | O | 2.77 | Y132 | OH | Hydrogen bond |
| **4** | P29 | O | 3.01 | R199 | NE | Hydrogen bond |
| **5** | R119 | NH2 | 3.26 | Y28 | OH | Hydrogen bond |
| **6** | N150 | ND2 | 3.36 | R199 | O | Hydrogen bond |
| **7** | D156 | OD2 | 3.31 | T240 | OG1 | Hydrogen bond |
| **8** | D156 | OD2 | 3.81 | T240 | N | Hydrogen bond |
| **9** | K198 | O | 2.60 | Q228 | NE2 | Hydrogen bond |
| **10** | R199 | O | 2.92 | N150 | ND2 | Hydrogen bond |
| **11** | R199 | NE | 3.23 | P29 | O | Hydrogen bond |
| **12** | V202 | O | 3.01 | V231 | N | Hydrogen bond |
| **13** | R204 | NH2 | 2.60 | P242 | OXT | Hydrogen bond |
| **14** | R204 | N | 2.84 | V231 | O | Hydrogen bond |
| **15** | R204 | O | 2.90 | V233 | N | Hydrogen bond |
| **16** | R204 | NH1 | 3.06 | D232 | OD1 | Hydrogen bond |
| **17** | R204 | NH1 | 3.47 | A241 | O | Hydrogen bond |
| **18** | R206 | N | 3.62 | V233 | O | Hydrogen bond |
| **19** | H227 | O | 3.12 | H227 | NE2 | Hydrogen bond |
| **20** | H227 | NE2 | 3.13 | H227 | O | Hydrogen bond |
| **21** | V231 | O | 2.82 | R204 | N | Hydrogen bond |
| **22** | V231 | N | 2.98 | V202 | O | Hydrogen bond |
| **23** | D232 | OD1 | 2.87 | R204 | NE | Hydrogen bond |
| **24** | D232 | OD2 | 2.96 | R204 | NH2 | Hydrogen bond |
| **25** | V233 | N | 2.81 | R204 | O | Hydrogen bond |
| **26** | V233 | O | 2.95 | R206 | N | Hydrogen bond |
| **27** | D236 | OD2 | 2.71 | Y210 | N | Hydrogen bond |
| **28** | D236 | OD2 | 2.91 | A208 | N | Hydrogen bond |
| **29** | D236 | OD1 | 2.95 | R211 | N | Hydrogen bond |
| **30** | D236 | N | 3.29 | R206 | O | Hydrogen bond |
| **31** | T240 | OG1 | 2.55 | D156 | OD1 | Hydrogen bond |
| **32** | T240 | OG1 | 2.65 | Y186 | OH | Hydrogen bond |
| **33** | T240 | N | 2.89 | D156 | OD2 | Hydrogen bond |
| **34** | A241 | O | 2.82 | R204 | NH1 | Hydrogen bond |
| **35** | A241 | O | 2.84 | R204 | NH2 | Hydrogen bond |
| **36** | A241 | N | 3.70 | Y186 | OH | Hydrogen bond |
| **37** | R204 | NH1 | 3.06 | D232 | OD1 | Salt bridge |
| **38** | R204 | NE | 3.52 | D232 | OD1 | Salt bridge |
| **39** | R204 | NH1 | 3.79 | D232 | OD2 | Salt bridge |
| **40** | D232 | OD1 | 2.87 | R204 | NE | Salt bridge |
| **41** | D232 | OD2 | 2.96 | R204 | NH2 | Salt bridge |
| **42** | D232 | OD1 | 3.40 | R204 | NH2 | Salt bridge |
| **43** | D232 | OD2 | 3.94 | R204 | NE | Salt bridge |

**S3 Table. Effect of HPLC conditions on the retention times of peaks in dAK assays**. The table shows the retention times of peaks in dAK assays with deoxyadenosine analogues using isocratic conditions with a binary mixture of different ratios of solution A and B (listed as A percentage). Peaks that are independent of the A percentage are marked in grey and only listed for the 75% A protocol. Protocols with low A percentage (7 or 20%) also include elution and equilibration steps as described below.

| | **HPLC protocols indicated by their mobile phase percentage of A** | | | | | |
|---|---|---|---|---|---|---|
| **Peak** | **7A** | **20A** | **62A** | **68A** | **75A** | **100A** |
| Ara-A | *Buffer-independent peak* | | | | 2.39 | |
| dAdo | *Buffer-independent peak* | | | | 2.77 | |
| FANA-A | *Buffer-independent peak* | | | | 3.33 | |
| F-Ara-A | *Buffer-independent peak* | | | | 3.99 | |
| F-dAdo | *Buffer-independent peak* | | | | 4.83 | |
| DTT | *Buffer-independent peak* | | | | 7.70 | |
| Cladribine | *Buffer-independent peak* | | | | 7.89 | |
| Clofarabine | *Buffer-independent peak* | | | | 11.69 | |
| AMP | 9.01 | 4.46 | | | 2.68 | |
| Ara-AMP | 9.65 | 4.67 | | | 2.72 | |
| dAMP | 12.59 | 5.97 | | | 3.23 | |
| FANA-AMP | 21.35 | 9.43 | | 5.11 | 4.53 | |
| F-Ara-AMP | 25.14 | 11.54 | | | 5.81 | |
| F-dAMP | 35.60 | 16.14 | | 8.43 | 7.74 | |
| Cladribine-MP | | 31.33 | 16.26 | | 14.45 | |
| Clofarabine-MP | | | | 29.46 | 25.38 | 22.98 |
| ADP | | 15.23 | 5.54 | 4.52 | 4.17 | |
| ATP | | (75.41)* | 10.83 | 9.53 | 8.23 | 6.48 |
| Minor peak 1 | | | 17.83 | 15.43 | 13.35 | |
| Minor peak 2 | | | 23.05 | 18.70 | 15.74 | |
| Minor peak 3 | | | 48.40 | 38.81 | 32.40 | 19.42 |
| *Analysis time* | *47*** | *43*** | *24.2*** | *19.4*** | *33* | *24* |

*Retention time if the isocratic conditions would be continued.

**Maximal analysis time if all listed assay products are studied. It includes the elution-equilibration procedure for the 7A and 20A protocols and is set to half the retention time of Minor peak 3 in the 62A and 68A protocols.

**75A and related HPLC protocols.** The 75A HPLC protocol was used for dAK assays with most deoxyadenosine analogues when tested at 200 µM (F-Ara-A, F-dAdo cladribine and clofarabine), both when assayed alone and in combination with deoxyadenosine. The samples were in these cases diluted 100 times before analysis, and the listed minor peaks are then barely visible. With more concentrated samples (5-20-fold dilutions), it was necessary to decrease the A percentage to 68% for F-dAdo and 62% for cladribine to avoid interference with the broad ATP peak and the minor peaks. The 100A protocol was used for $K_M$-$V_{max}$ measurements with clofarabine to reduce the analysis time.

**7A and 20A.** These protocols were originally optimized for dAK assays with Ara-A, FANA-A with and without deoxyadenosine as competitor, but they are also usable for assays with most other deoxyadenosine analogies (products listed above). Both protocols require an additional 10-min step with 90% A to remove late-eluting peaks followed by a 10-min equilibration step to initial conditions. The 20A protocol gives a faster elution time for each assay product, but the 7A protocol has the advantage of a better separation between Ara-AMP and AMP.

**S4 Table. Cryo-EM data collection statistics of *G. intestinalis* dAK.**

| **Data collection** | **1** | **2** |
|---|---|---|
| Microscope | Titan Krios | Titan Krios |
| Detector | K2 (Gatan) | Falcon 4i |
| Energy filter slit (eV) | 20 | 20 |
| Voltage (kV) | 300 | 300 |
| Magnification | 215000 | 215000 |
| Objective Aperture (µm) | 100 | 100 |
| Specimen pixel size (Å/px) | 0.63 | 0.58 |
| Total dose per movie ($e^-$/Å$^2$) | 59.5 | 60 |
| Defocus range (µm) | -1.2 to -2.4 | -0.5 to -3.5 |
| Spherical aberration (nm) | 2.7 | 2.7 |
| Number of micrographs | 2534 | 3420 |

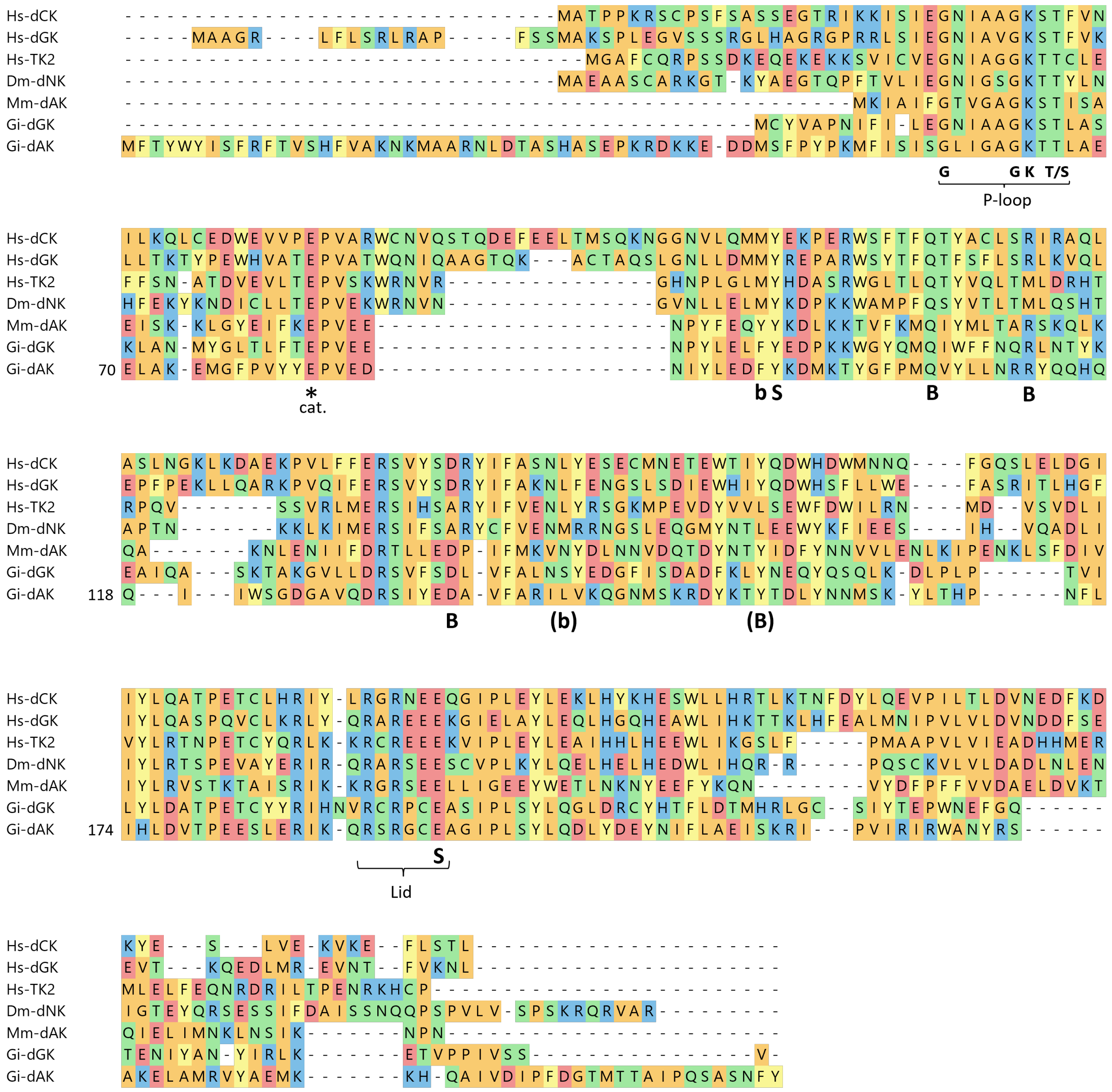

**S1 Fig. Amino acid sequence alignment of *G. intestinalis* dAK with other non-TK1-like dNKs.** The most striking difference of the *G. intestinalis* dAK compared to the other sequences is its longer termini, particularly the N-terminus. Conserved features are the P-loop for binding the phosphate donor, the catalytic base (labeled *) and the lid domain. Based on the published *Mycoplasma mycoides* dAK structure, active site residues in contact with the substrate are labeled with B and S depending on if they bind the nucleobase or sugar moiety. Non-capital letters mean that the residues affect the binding of the substrate indirectly and brackets mean that they are not conserved in the *G. intestinalis* dAK.

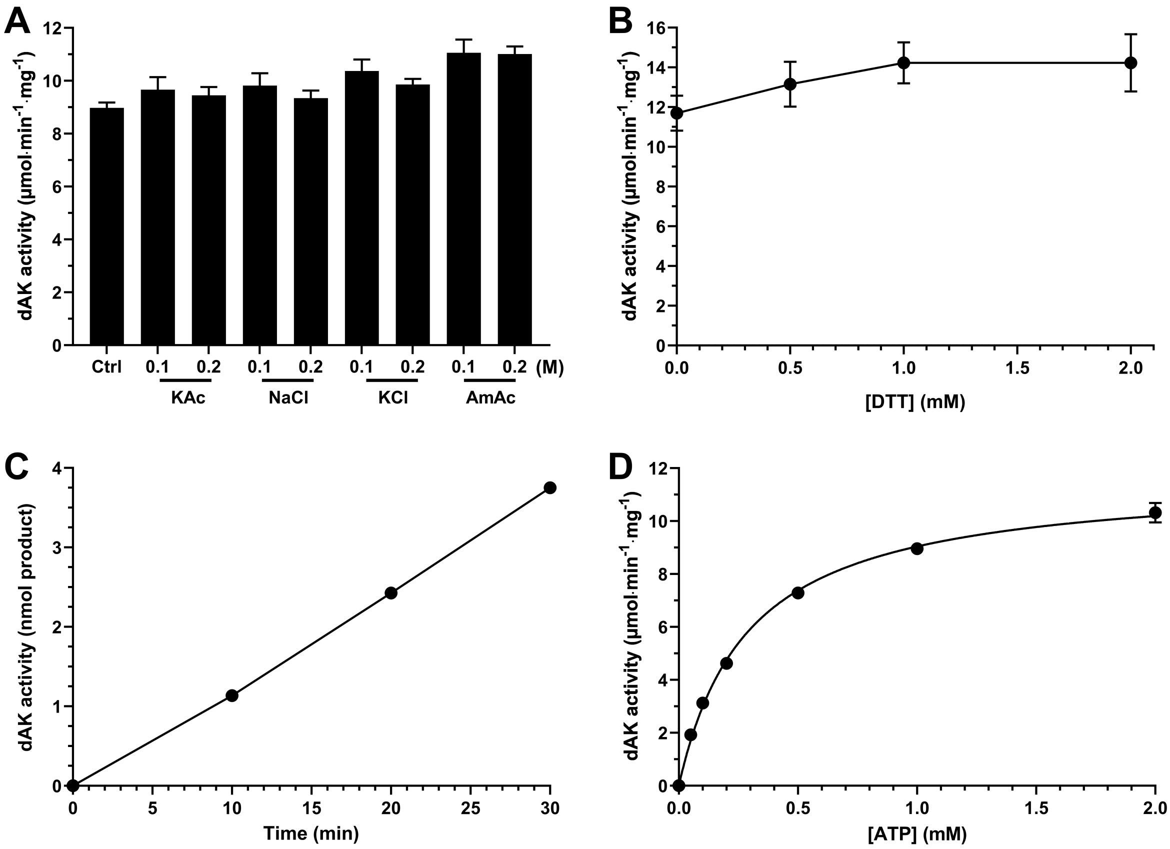

**S2 Fig. Development of enzyme assay conditions for *G. intestinalis* dAK**. (A) Effect of different chloride and acetate salts on enzyme activity. (B) Effect of various concentrations of DTT on enzyme activity. (C) Testing the linearity of the enzyme activity over time. (D) Effect of the ATP concentration on enzyme activity. The enzyme assays were generally performed with 10 ng dAK, 200 µM deoxyadenosine, 2 mM ATP (variable in D), 5 mM $MgCl_2$, 0.5 mM DTT (variable in B) 100 mM potassium acetate (variable salts in A) and 50 mM Tris-HCl pH 7.5 at 37°C for 30 min (variable in C). A, B and D represent the average of three independent experiments with standard errors indicated. A general conclusion was that the tested salts had no major effect on enzyme activity when used up to 200 mM, DTT was slightly stimulatory, the activity was linear over 30 min and that a concentration of 2 mM ATP was sufficient to make it the non-limiting substrate under standard assay conditions. The $K_M$ of ATP calculated from D was 0.29 ± 0.03 mM with the standard error based on the results from the individual experiments.

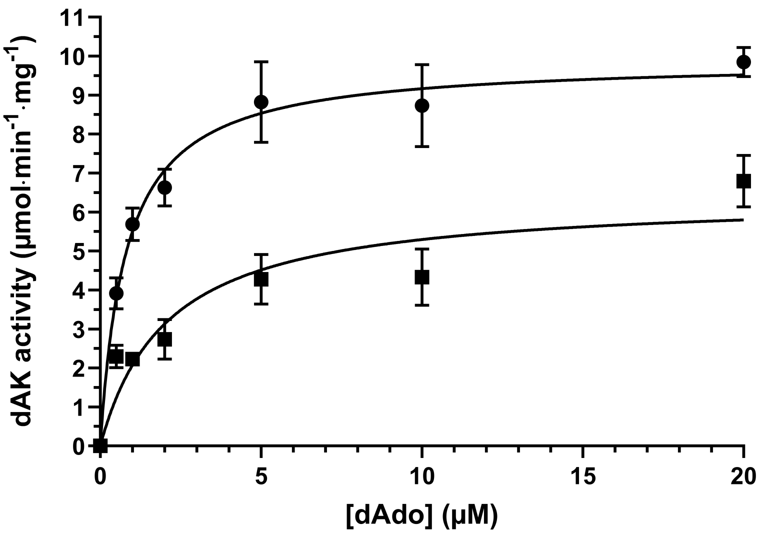

**S3 Fig. *G. intestinalis* dAK is subject to mixed model inhibition by dATP**. The enzyme assay was performed in the absence (●) or presence (■) of 1 µM dATP and the curves were fitted to a mixed inhibition model by GraphPad Prism 10.0.2. The experimental parameters obtained by the experiment were the following: $K_i$= 0.34 ± 0.05 µM, $K_M$=0.80 ± 0.10 µM, $V_{max}$=9.9 ± 0.9 µmmol·min$^{-1}$·mg$^{-1}$ and α=5.5 ± 0.9 with standard errors calculated from the variation between the individual three experiments. The inhibition model chosen gave the best fit to the data points and it was also supported by that the $K_M$ was increased in the presence of dATP ($K_{M,app}$=2.0 ± 0.2 µM) and the $V_{max}$ was decreased ($V_{max,app}$=6.4 ± 0.9 µmmol·min$^{-1}$·mg$^{-1}$). The graph represents the average of three independent experiments with standard errors indicated.

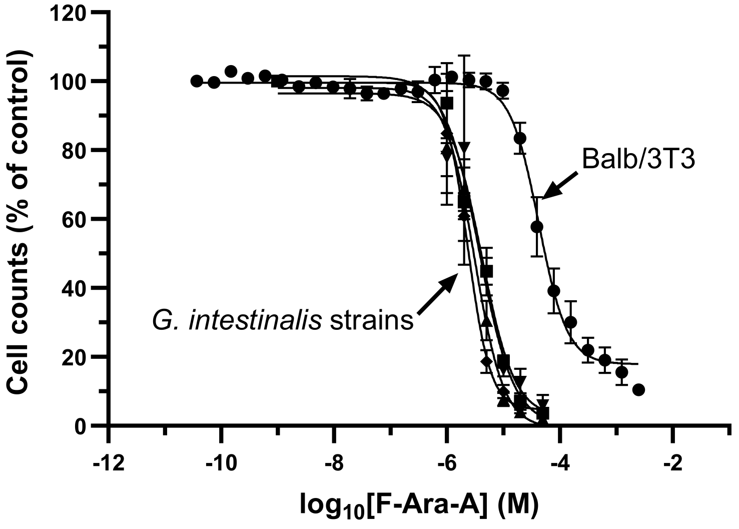

**S4 Fig. F-Ara-A selectively inhibits *G. intestinalis* proliferation**. The figure illustrates that *G. intestinalis* is more sensitive than the mammalian reference cell line to F-Ara-A and that the parasite strains are equally affected by the drug regardless of if they are metronidazole-resistant or not. The cells used in the experiments are mammalian Balb/3T3 fibroblasts (●), the *G. intestinalis* parent strain WB (■), the metronidazole-resistant strains M1 (▲), the metronidazole-resistant strain M2 (▼), and the resistant revertant strain M1-NR (◆). The graphs represent the average of 3 or more experiments with standard errors and the data is fitted to a log [inhibitor] vs response (variable slope) curve by GraphPad Prism 10.1.0.

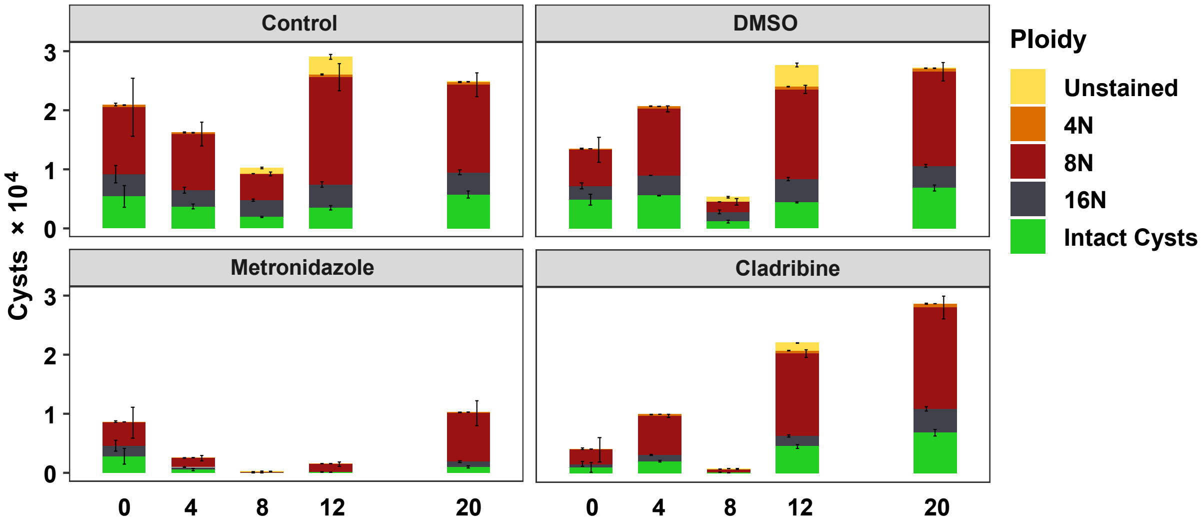

**S5 Fig. Cladribine inhibits early stages of *G. intestinalis* encystation**. The results show the ploidy of the cells after 28 hours of incubation in encystation medium with the indicated drugs added in the beginning of the incubation or at indicated timepoints during the encystation process. The drugs were solubilized in DMSO prior to analysis. Control experiments were performed in a similar manner but with no liquid or DMSO added to the samples instead of drug. Cladribine treatment needed to be initiated within the first 12 hours to have a significant effect and then the main features were a marked reduction in the number of cells (if started within the first 8 h) and a delayed progression into 16N cells (also observable if given at the 12 h timepoint). The control cells were much less dependent on when they were treated except for the 8-h timepoint when the cells are going through a membrane reorganization process that makes them more sensitive to disturbances.

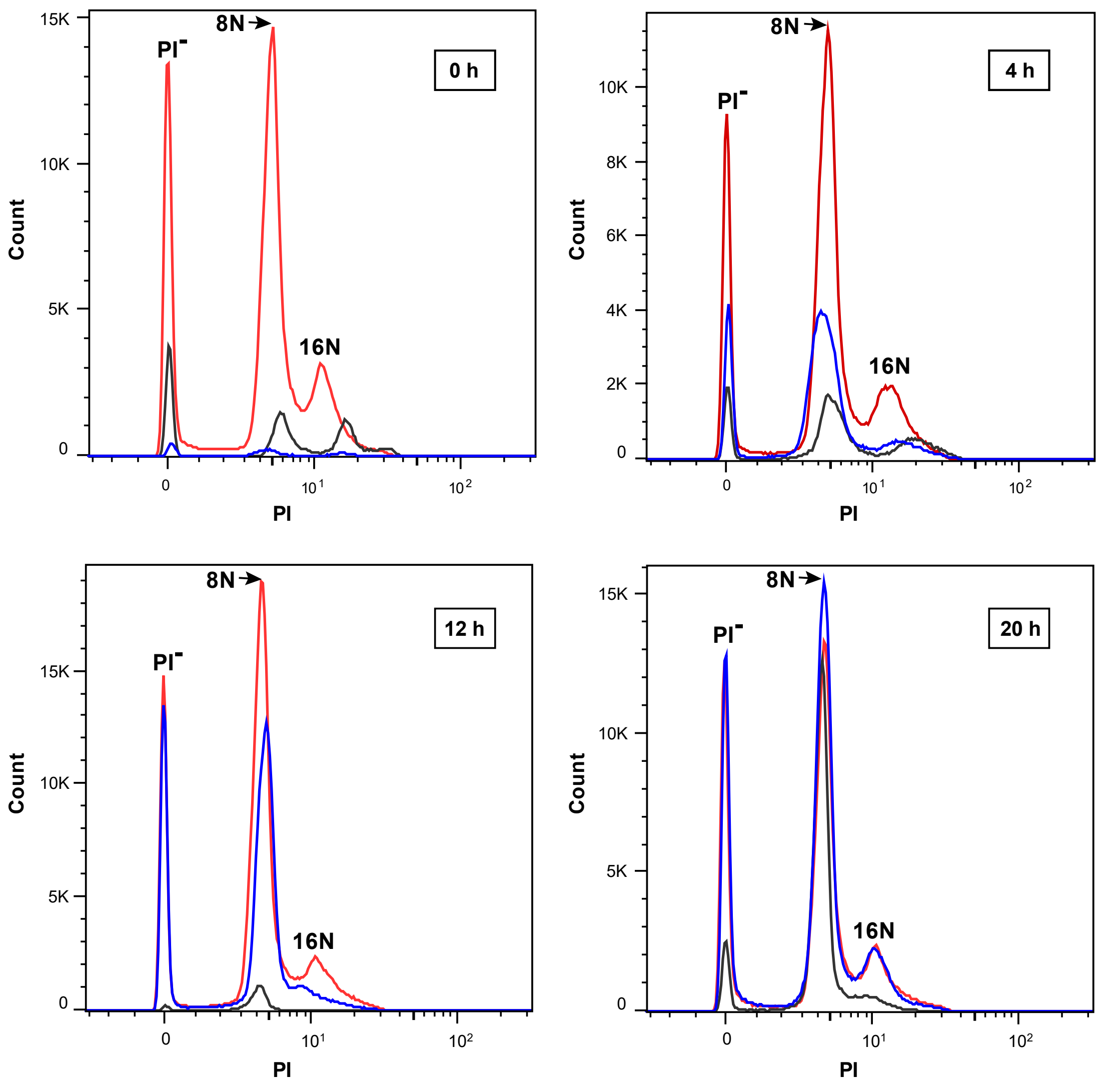

**S6 Fig. PI profiles from the encystation experiments**. Similar type of experiment as in S5 Fig but showing the PI profile. The color code indicates control cells (red), metronidazole-treated cells (grey) and cladribine-treated cells (blue). Cladribine has a strong effect on the total number of cells if given early in the process and on the number of 16N cells if given at 12 h, but no obvious effect if given late in the process (20 h).

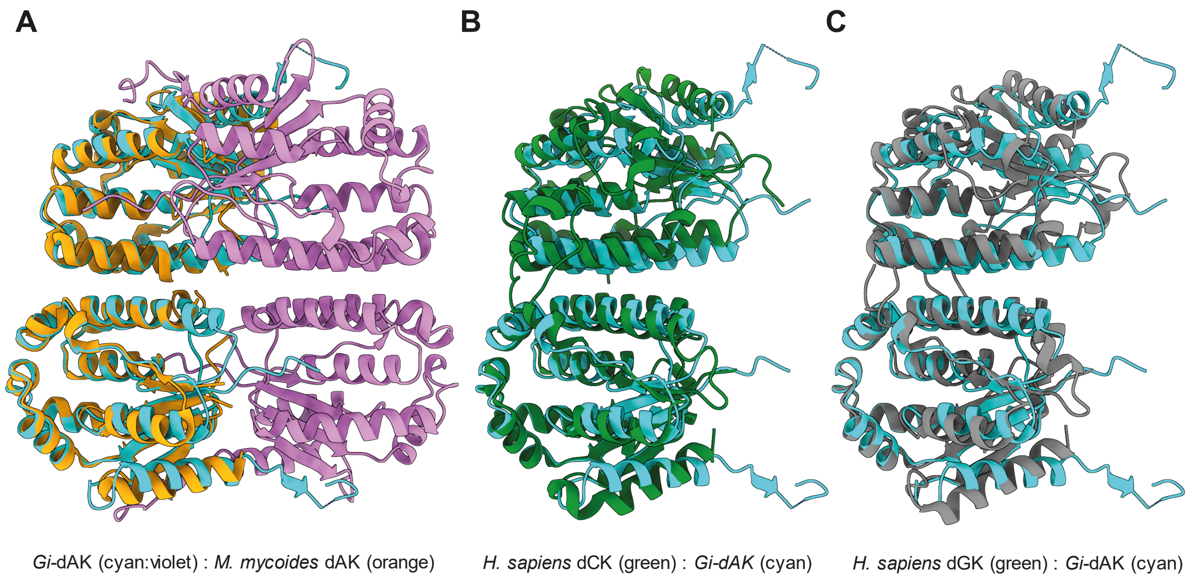

**S7 Fig. Comparison of the *G. intestinalis* dAK structure with other non-TK1-like dNKs**. Although the structural fold of the polypeptides is similar in the three examples, *G. intestinalis* dAK (labeled *Gi*-dAK) sticks out by being tetrameric. Note that it is only in A that the whole tetramer is shown, whereas B-C is focused on the dimer part to highlight the homology. (A) Tetrameric structure of *G. intestinalis* dAK (cyan/magenta) superimposed onto the structure of *M. mycoides* dAK (orange, PDBID: 2JAQ). (B) *G. intestinalis* dAK dimer (cyan) superimposed onto *H. sapiens* dCK (green, PDBID: 2ZI3). (C) *G. intestinalis* dAK dimer (cyan) superimposed onto *H. sapiens* dGK (grey, PDBID: 2OCP). Protein structures are shown as cartoon representation.

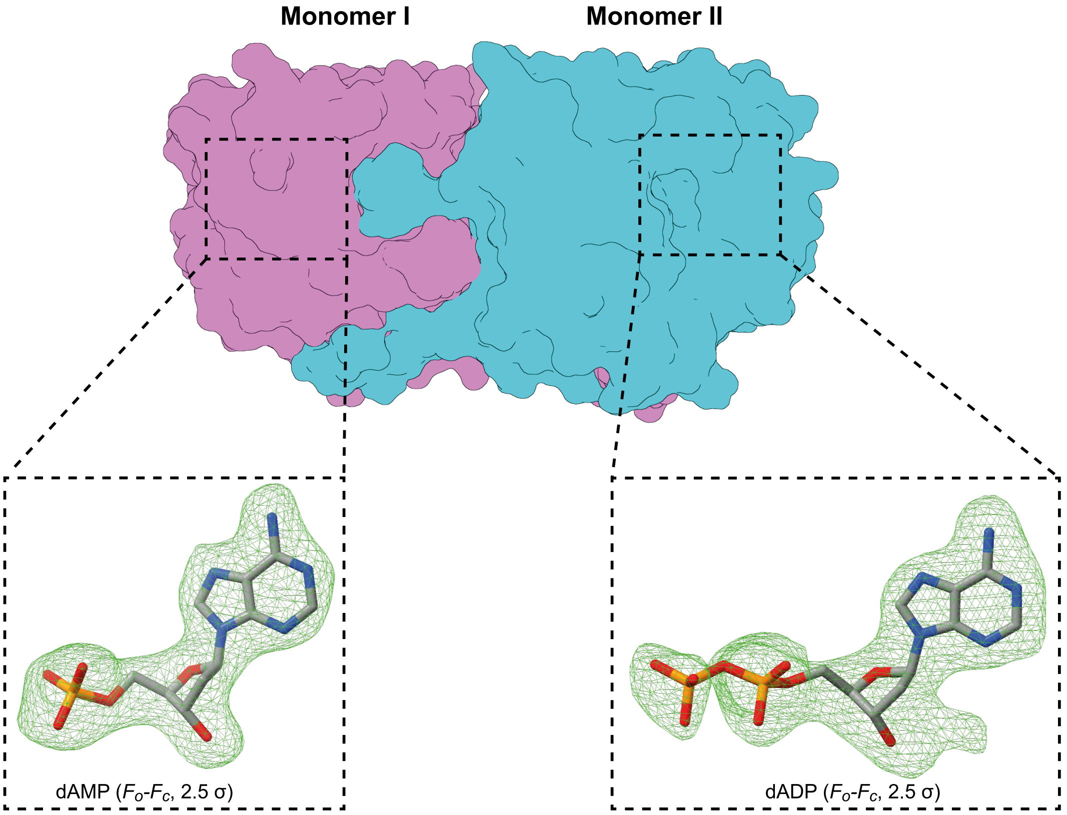

**S8 Fig. Unbiased $F_o$-$F_c$ map showing the placement of dADP and dAMP in *G. intestinalis* dAK**. The crystallographic dimer of *G. intestinalis* dAK revealed that two ligands were co-purified during protein purification. Monomer I (magenta) contains density for a dAMP molecule, and monomer II (cyan) contains density belonging to a dADP molecule, with a partial (0.5) occupancy for the β-phosphate. dAMP and dADP are shown as grey sticks and the unbiased $F_o$-$F_c$ density map as a green mesh contoured to 2.5 s.

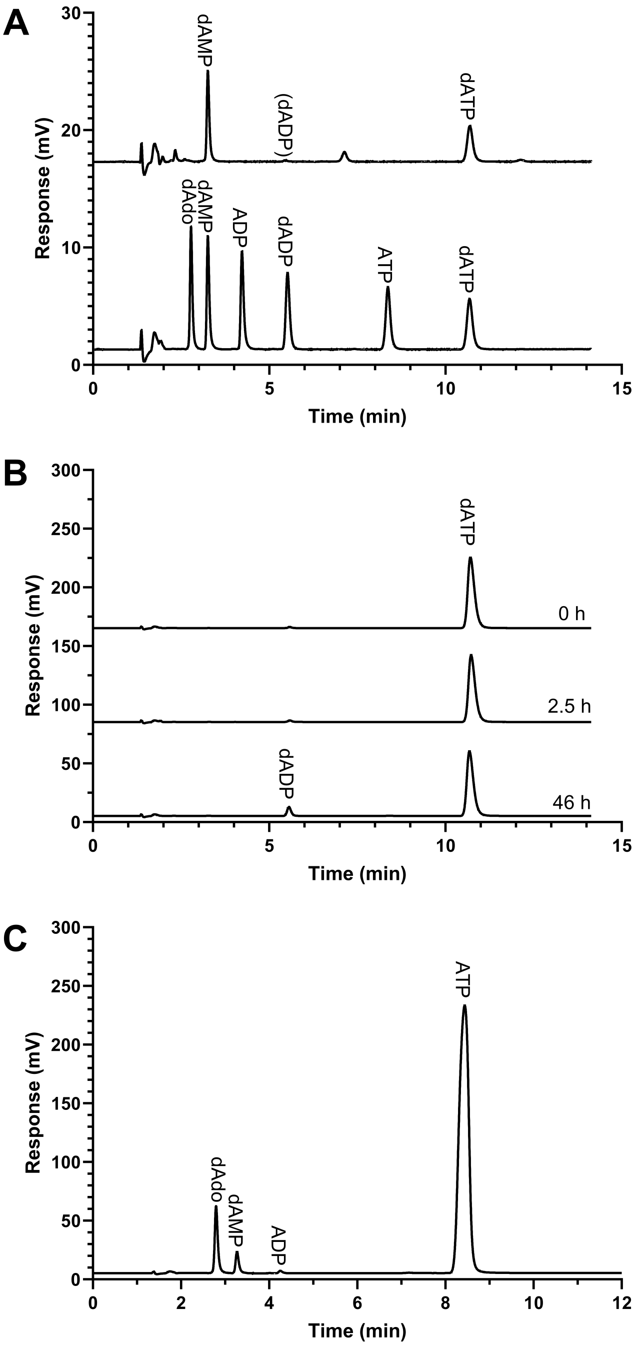

**S9 Fig. HPLC analysis of nucleotides in the *G. intestinalis* dAK preparation**. (A) The dAK preparation contains dAMP and dATP. (B) *G. intestinalis* dAK incubated with 200 µM dATP at room temperature over 46 h shows that dATP is slowly converted to dADP. (C) Analys of dAK in a regular enzyme assay shows that this activity is much higher than the dATP-converting activity, with the latter possibly being the result of a minor impurity. The regular enzyme assay was performed with 10 ng dAK over 30 min (C) as compared to 100 ng dAK and 46 hours used in the dATP-dephosphorylation assay (B).

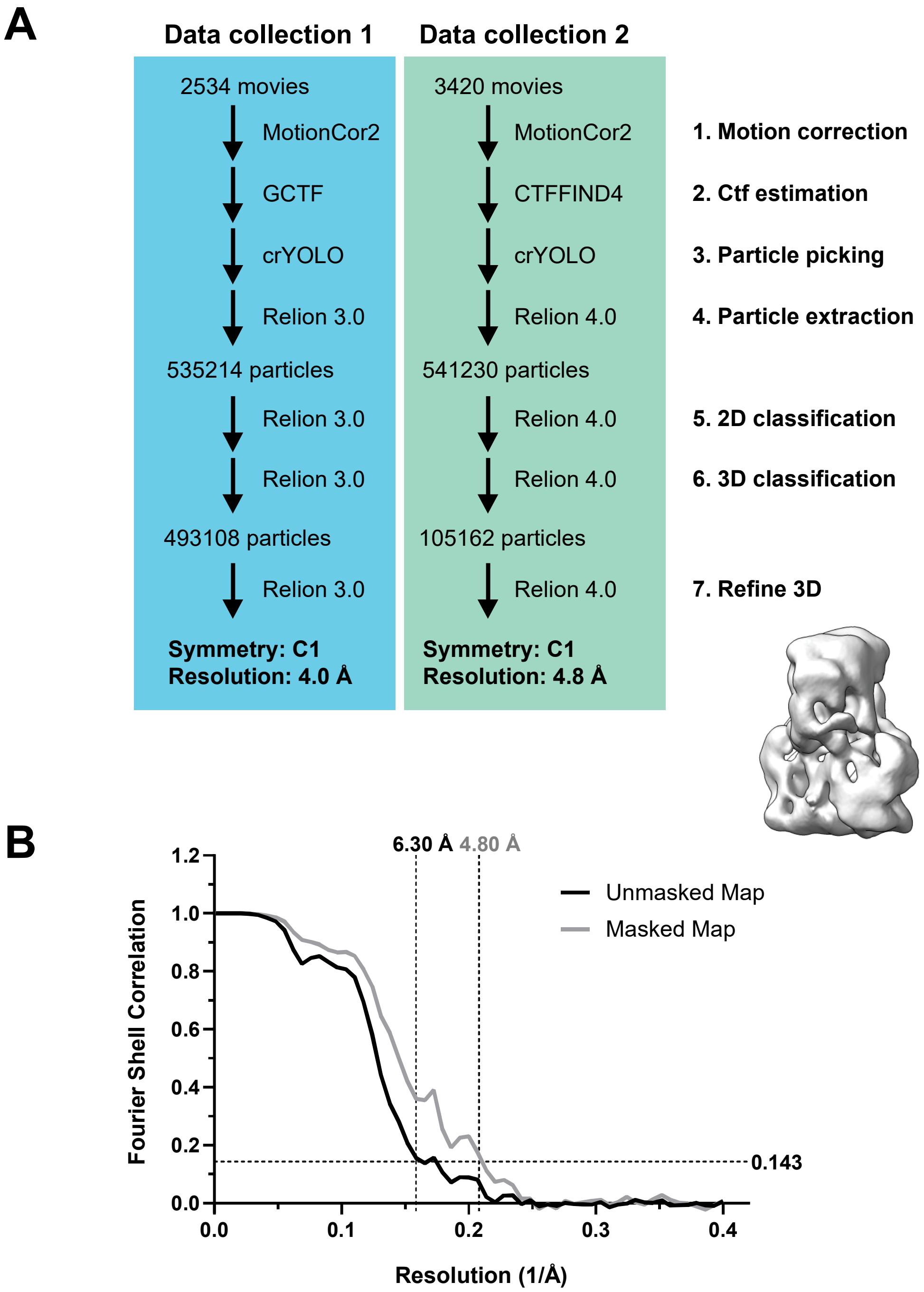

**S10 Fig. Cryo-EM methodology**. (A) Cryo-EM data collection flowchart. (B) Graphs showing the Gold standard Fourier-Shell-Correlation (FSC) for independently refined half sets vs. resolution [1/Å] of the unmasked and masked volumes for data collection 2. FSC at 0.143 is indicated as a black dashed line. Data collection 1 generated higher quality 2D class averages and is used as the source of the images in Fig. 5B. Data collection 2 generated a 3D map that looked slightly less anisotropic and is the source of the 3D map shown in Fig. 5C.

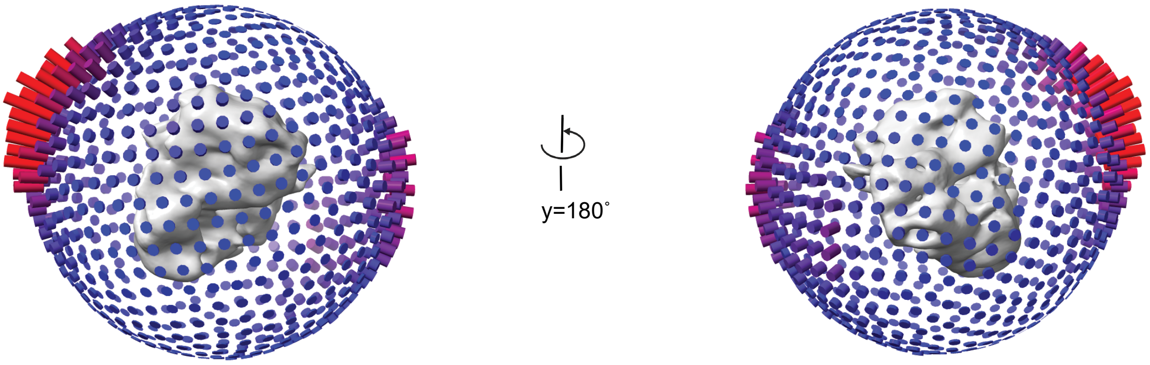

**S11 Fig. *G. intestinalis* dAK particles on grids have a strong preferred orientation**. A frequency diagram showing the number of views from a particular direction in the dataset as the length of each bar surrounding the 3D model. The longer the red bar, the more particles present whereas purple and blue show underrepresented views with fewer particles. The image thus indicates that there are many views that are missing to build an accurate 3D model of dAK from the cryo-EM data.
